# Fluorescent and bioluminescent calcium indicators with tuneable colors and affinities

**DOI:** 10.1101/2021.11.13.468356

**Authors:** Nicole Mertes, Marvin Busch, Magnus-Carsten Huppertz, Christina Nicole Hacker, Clara-Marie Gürth, Stefanie Kühn, Julien Hiblot, Birgit Koch, Kai Johnsson

**Affiliations:** Department of Chemical Biology, Max Planck Institute for Medical Research, Jahnstrasse 29, 69120 Heidelberg, Germany; Department of Optical Nanoscopy, Max Planck Institute for Medical Research, Jahnstrasse 29, 69120 Heidelberg, Germany; Institute of Chemical Sciences and Engineering, École Polytechnique Fédérale de Lausanne (EPFL), 1015 Lausanne, Switzerland

**Keywords:** Fluorescence, Bioluminescence, Calcium Indicator

## Abstract

We introduce a family of bright, rhodamine-based calcium indicators with tuneable affinities and colors. The indicators can be specifically localized to different cellular compartments and are compatible with both fluorescence and bioluminescence readouts through conjugation to HaloTag fusion proteins. Importantly, their increase in fluorescence upon localization enables no-wash live-cell imaging, which greatly facilitates their use in biological assays. Applications as fluorescent indicators in rat hippocampal neurons include the detection of single action potentials and of calcium fluxes in the endoplasmic reticulum (ER). Applications as bioluminescent indicators include the recording of the pharmacological modulation of nuclear calcium in high-throughput-compatible assays. The versatility and remarkable ease of use of these indicators make them powerful tools for bioimaging and bioassays.

**Graphical abstract:** 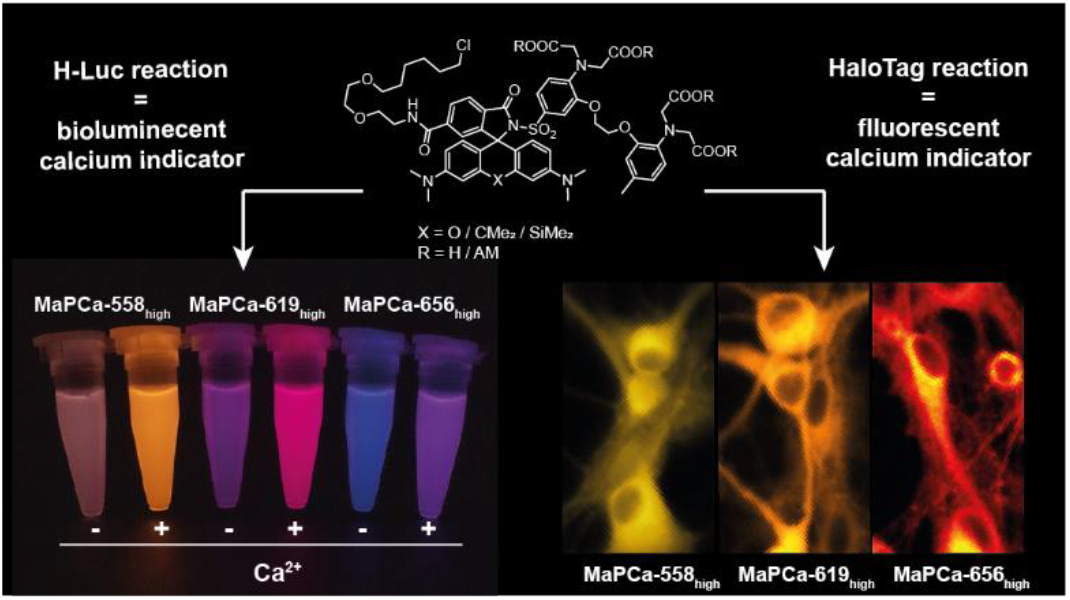

## Introduction

The second messenger calcium is involved in a plethora of signaling pathways and biochemical processes.^1^ The elucidation of its function in cellular processes has become possible largely through the development of calcium indicators.^2–4^ While early development focused on synthetic calcium indicators, genetically encoded calcium indicators (GECIs) have now become the gold standard. The main reason for this is that GECIs can be genetically targeted to specific cellular populations and subcellular localizations, whereas the cellular uptake of synthetic calcium indicators lacks selectivity and is often inefficient. However, GECIs possess lower brightness, slower response kinetics and a limited color range (especially in the far-red in comparison to synthetic indicators).^5,6^ These limitations are of particular concern when highly localized areas, such as micro- and even nanodomains are investigated, and more demanding microscopy techniques are used.^7–9^ A possibility to combine the brightness, response kinetics and spectral range of synthetic fluorescent indicators with the targetability of GECIs is the use of self-labeling protein tags such as SNAP-tag and HaloTag.^10,11^ Self-labeling proteins form a covalent bond to a specific substrate and through this enable precise localization of synthetic molecules to proteins of interest (POI). This approach has been used to create a number of localizable synthetic calcium indicators, *e.g.* BG3-Indo-1,^12^ BOCA-1-BG^13^ or RhoCa-Halo^14^ and the far-red indicator JF_646_-BAPTA.^5,15^ However, these probes have limited cell permeability and solubility, and furthermore require washing steps to remove unreacted probes, greatly limiting their applicability.^13,14^ The use of bright synthetic fluorophores for calcium sensing was enabled developing chemogenetic sensors in which the protein-based calcium-sensing domain Calmodulin (CaM) interacts with an environmentally sensitive dye (*e.g.* rHCaMP or HaloCaMP).^16,17^ However, based on the same calcium sensing domain as most GECIs, they suffer from relatively slow response kinetics.^16^ Furthermore, there is currently no localizable synthetic far-red calcium indicator with a suitable calcium affinity for calcium-rich areas like the endoplasmic reticulum (ER) or calcium microdomains.^18,19^ Here we present MaPCa dyes, a family of highly permeable calcium indicators with different colors and calcium affinities that can be coupled to HaloTag. As the reaction with HaloTag shifts the fluorescent scaffold of the indicator from a non-fluorescent into a fluorescent configuration, these probes can be used without any washing steps to remove unbound probe.

## Results and discussion

### Design principle, synthesis and in vitro characterization of MaPCa dyes

The design of our calcium indicators is based on the recently introduced MaP dyes, in which the lactone-forming carboxylic acid of a rhodamine is replaced with an amide attached to an electron withdrawing group (*e.g.* sulfonamides).^20,21^ This results in dyes that preferentially exist as a non-fluorescent spirolactam in solution, but shift to an open, fluorescent state upon binding to HaloTag, enabling no-wash imaging with low background. We envisioned designing fluorogenic calcium indicators by attaching a calcium chelator such as BAPTA (1,2-bis(o-aminophenoxy)ethane-*N,N,N′,N*′-tetraacetic acid) through a benzene sulfonamide to the ortho-carboxylate of rhodamines and a chloroalkane (CA) through a carboxylate at the 6-position of the benzyl-ring (Fig. 1). BAPTA would be thereby positioned in close proximity to the rhodamine core, which is an important factor for effective PET-quenching of the rhodamine by the free chelator.^15,22^ Attachment of the CA via the 6-position of the benzyl-ring would enable HaloTag to shift the equilibrium from spirocyclization to the fluorescent, open form, thereby resulting in fluorogenicity. Furthermore, attachment of the CA via the 6-position would ensure a high labeling speed of the resulting HaloTag substrate.^23^

**Fig. 1:**
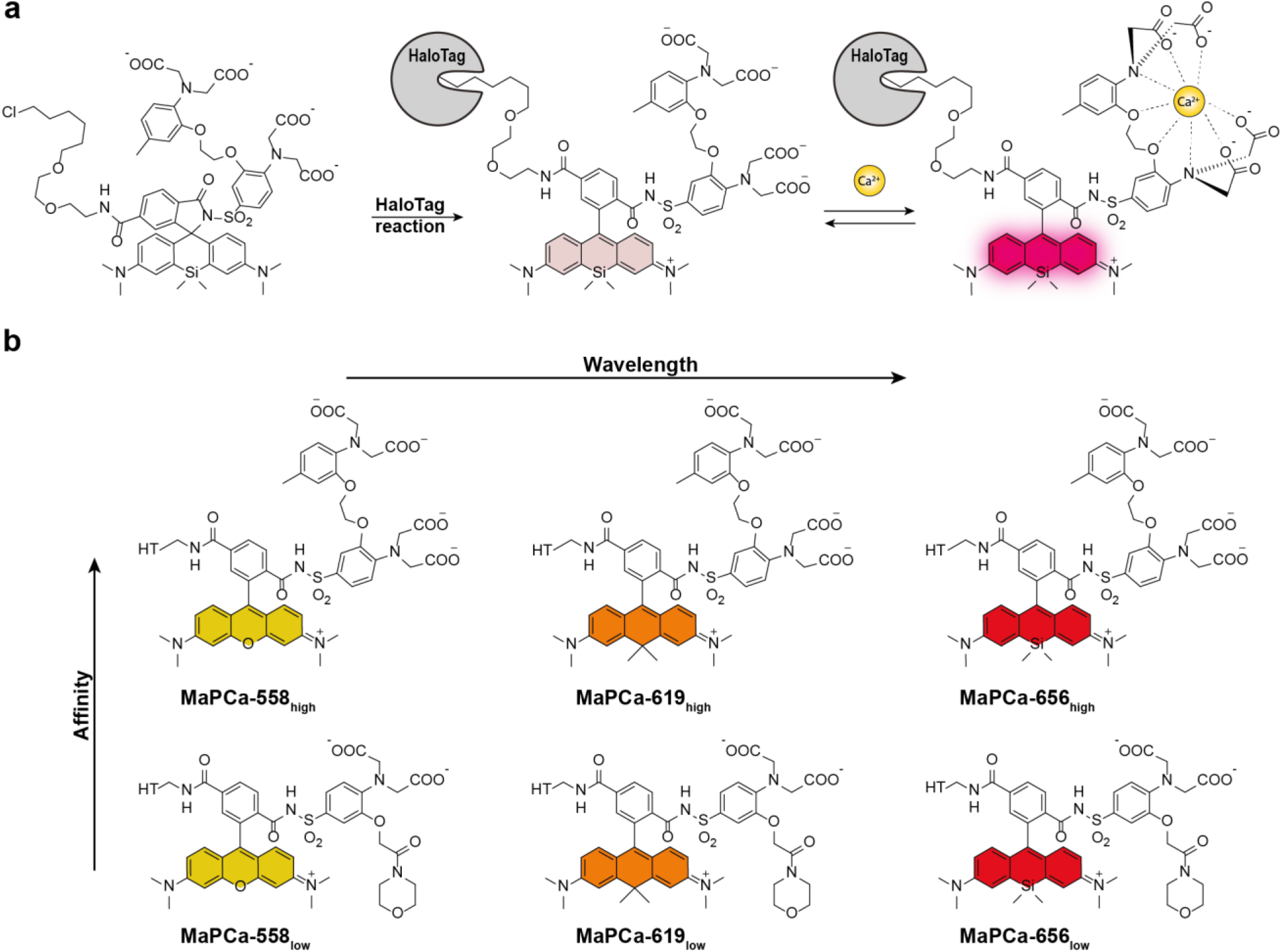
Schematic representation of the MaPCa dyes. a) Representation of the double-turn-on mechanism of MaPCa dyes. Example for MaPCa-656_high_. If not bound to the HaloTag, MaPCa dyes are in their colorless, spirocyclic form. Upon binding to HaloTag, they open to their zwitterionic form and hence become potentially fluorescent, but PET-quenched by the Ca^2+^-binding moiety. Only upon calcium binding full fluorescence is achieved. b) Overview of synthesized MaPCa dyes. HT= HaloTag-bound linker.

We set out to synthesize a set of such indicators based on the high-affinity calcium chelator BAPTA and the low-affinity chelator MOBHA (2-(2′-morpholino-2′-oxoethoxy)-*N,N*-bis(hydroxycarbonylmethyl) aniline)^24^ in combination with commercially available rhodamine-CA substrates TMR-CA, CPY-CA and SiR-CA, covering the spectrum from 550 to 650 nm (Fig. 2). In a first step, a sulfonamine was attached to the previously described BAPTA-ethylester^25^ **(01)** or MOBHA-ethylester **(02)** via chlorosulfonation followed by amination **(03,04)**. These two intermediates were then coupled to the commercially available rhodamine-CAs TMR-CA, CPY-CA and SiR-CA using activation by chlorosulfonic acid. The indicators were obtained as free acids after saponification with KOH (Fig. 2a). For *in cellulo* experiments, acetoxymethyl (AM) esters of the indicators were synthesized by prior transesterification of the chelator **(05,06)** and subsequent coupling to the fluorophore. The AM-esters serve to mask the carboxylic acids to ensure cell-permeability, but are cleaved inside the cell by endogenous esterases.^26^ We named these indicators MaPCa dyes (for Max-Planck-Calcium sensor), with a postfix expressing the absorption maxima in nm (TMR = 558; CPY = 619; SiR = 656) and the subscripts ‘*high’* or ‘*low’* for indicating the calcium affinity range. The AM-esters of the dyes are marked with an additional AM, in contrast to the saponified probes. It should be noted that this short and convergent synthetic scheme should enable the conversion of most rhodamine-CAs into calcium sensors in a single step.

**Fig. 2:**
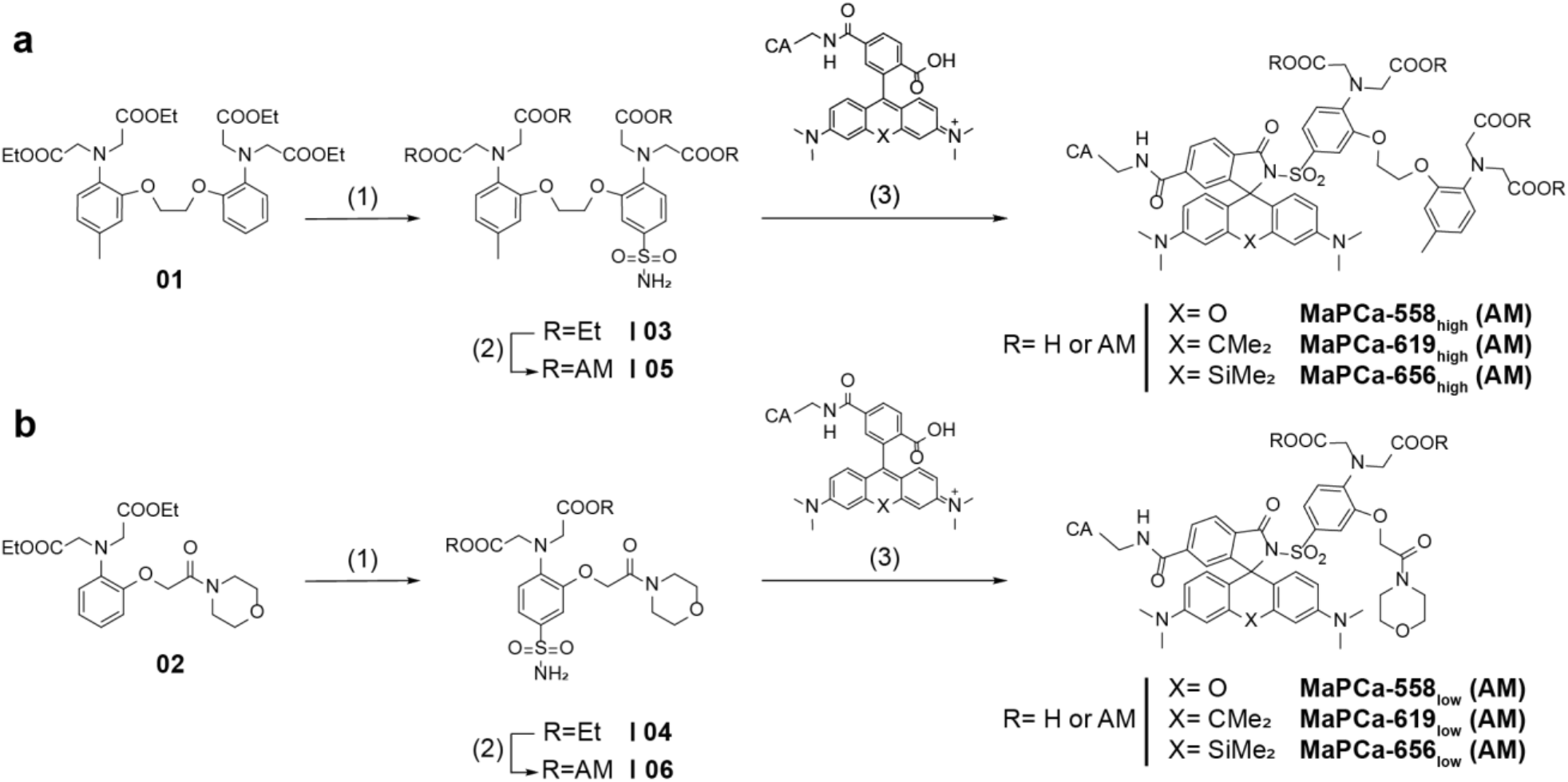
Synthetic pathway for the preparation of MaPCa dyes. The AM esters of the dyes are marked with an additional AM, in contrast to the saponified probe. (a) Synthetic route for MaPCa_high_ (1) i) HSO_3_Cl, SOCl_2_, DCM, 0°C-rt, 24 h ii) aq. NH_3_, EtOAc, rt, 75%; (2) this step was only performed for the AM-probes for the cellular experiments: i) DMAP, di-tert-butyl-dicarbonate, DCM, 35°C, 24 h ii) KOH, MeOH/THF, rt, 2 h iii) DIPEA, bromomethyl acetate, MeCN, rt, 48 h iv) TFA, DCM, rt, 2 h, 61%; (3) i) fluorophore preactivation with SOCl_2_, pyridine, DCM, rt – 60°C, 0.5 h ii) DIPEA, DMAP, 60°C, 1 h, 26-56%; the ethylester was subsequently saponified: KOH, MeOH/THF, rt, 8 h, 42-66%. (b) Synthetic route for MaPCa_low_ (1) i) HSO_3_Cl, SOCl_2_, DCM, 0°C-rt, 24 h ii) aq. NH_3_, EtOAc, rt, 44%; (2) this step was only performed for the AM-probes for the cellular experiments: i) DMAP, di-tert-butyl-dicarbonate, DCM/MeCN, 35°C, 43 h ii) KOH, MeOH/THF, rt, 5.5 h iii) DIPEA, bromomethyl acetate, MeCN, rt, 21 h iv) TFA, DCM, TIPS, rt, 0.5 h, 36%; (3) i) fluorophore preactivation with SOCl_2_, pyridine, DCM, rt – 60°C, 0.5 h ii) DIPEA, DMAP, 60°C, 3.5 h, 13-32%; the ethylester was subsequently saponified: KOH, MeOH/THF, rt, 5 h, 22-58%.

The MaPCa dyes calcium responsiveness was characterized *in vitro* in presence and absence of HaloTag measuring their fluorescence intensity at different free calcium concentrations (Fig. 3a, 3b, Supporting Fig. 3,4). As desired, all three high-affinity indicators showed a fluorogenic turn-on upon binding to HaloTag. However, while MaPCa-558_high_ was only slightly fluorogenic (1.3-fold), MaPCa-619_high_ and MaPCa-656_high_ showed a significant 7-fold and even 120-fold increase upon binding to HaloTag, respectively, in the calcium-bound state. The higher fluorogenicity of MaPCa-656_high_ can be rationalized considering the higher propensity of SiR derivatives to exist in the non-fluorescent spirocyclic form than the corresponding rhodamine and carborhodamine derivatives.^27^ The dyes possess a high brightness in the calcium-bound state (quantum yield >40%; extinction coefficient >80’000 M^−1^ cm^−1^) and display calcium-affinities in a suitable range for cytosolic measurements (K_D_(Ca^2+^): 410-580 nM) with turn-ons of around 6-fold upon calcium binding (Table 1, Supporting Table 1). The low-affinity indicators show similar fluorogenicities as the BAPTA-variants: the TMR-variant (MaPCa-558_low_) shows low fluorogenicity (1.4-fold) upon HaloTag binding, while MaPCa-619_low_ (28-fold) and MaPCa-656_low_ (208-fold) are highly fluorogenic. The calcium affinities of these dyes are in the range of 220-460 μM and they show a 7 to 11-fold turn-on upon calcium binding (Table 1, Supporting Table 1). The extinction coefficient of MaPCa-656_low_ is significantly lower than those of the other MaPCa-indicators, suggesting that it does not fully convert to the open state. Nevertheless, its brightness of ~15 mM^−1^ cm^−1^ is in the same order of magnitude than genetically encoded red-shifted indicators (brightness FR-GECO1c: 9.3 mM^−1^ cm^−1^).^28^

**Table 1:**
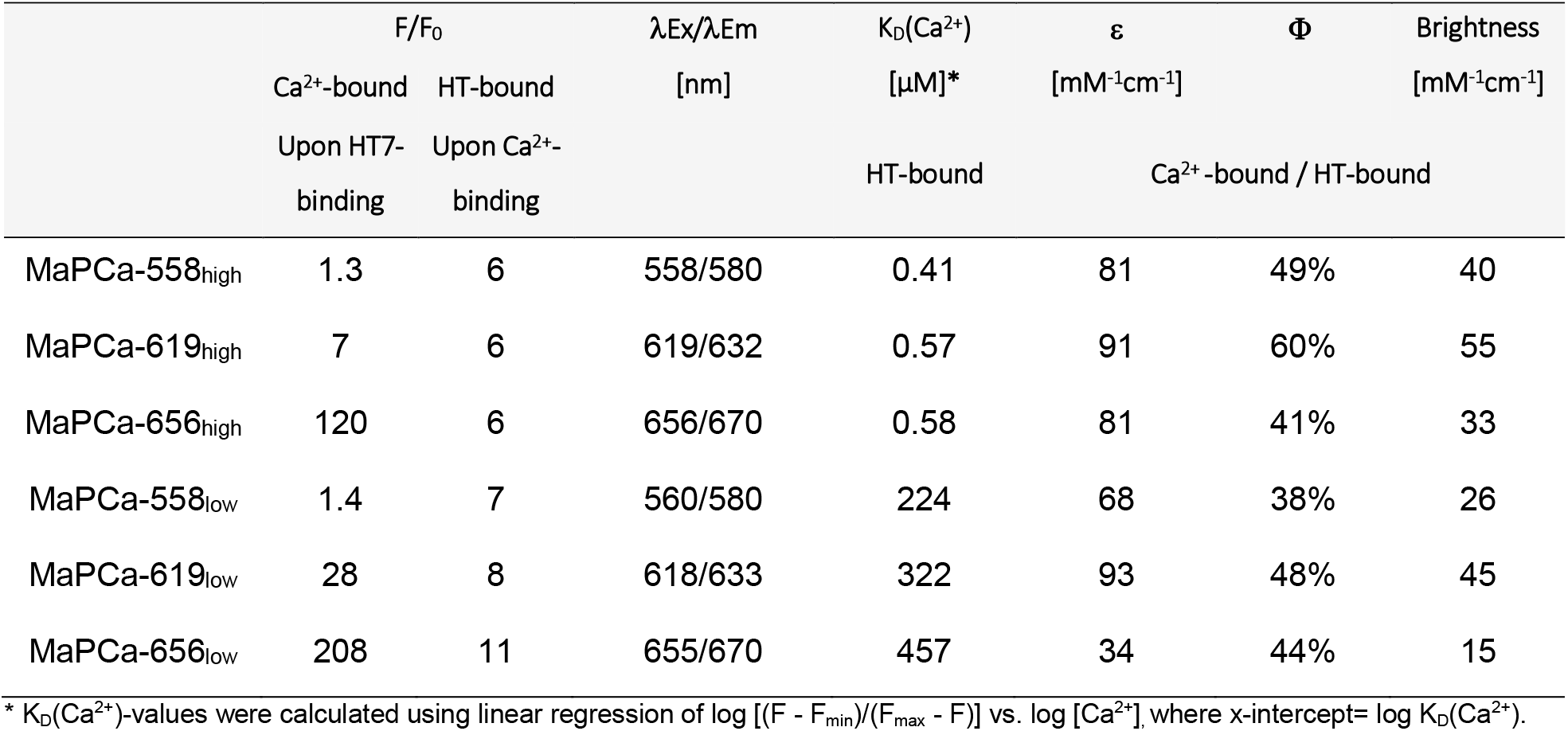
Photophysical properties of MaPCa dyes. HT= HaloTag.

**Fig. 3:**
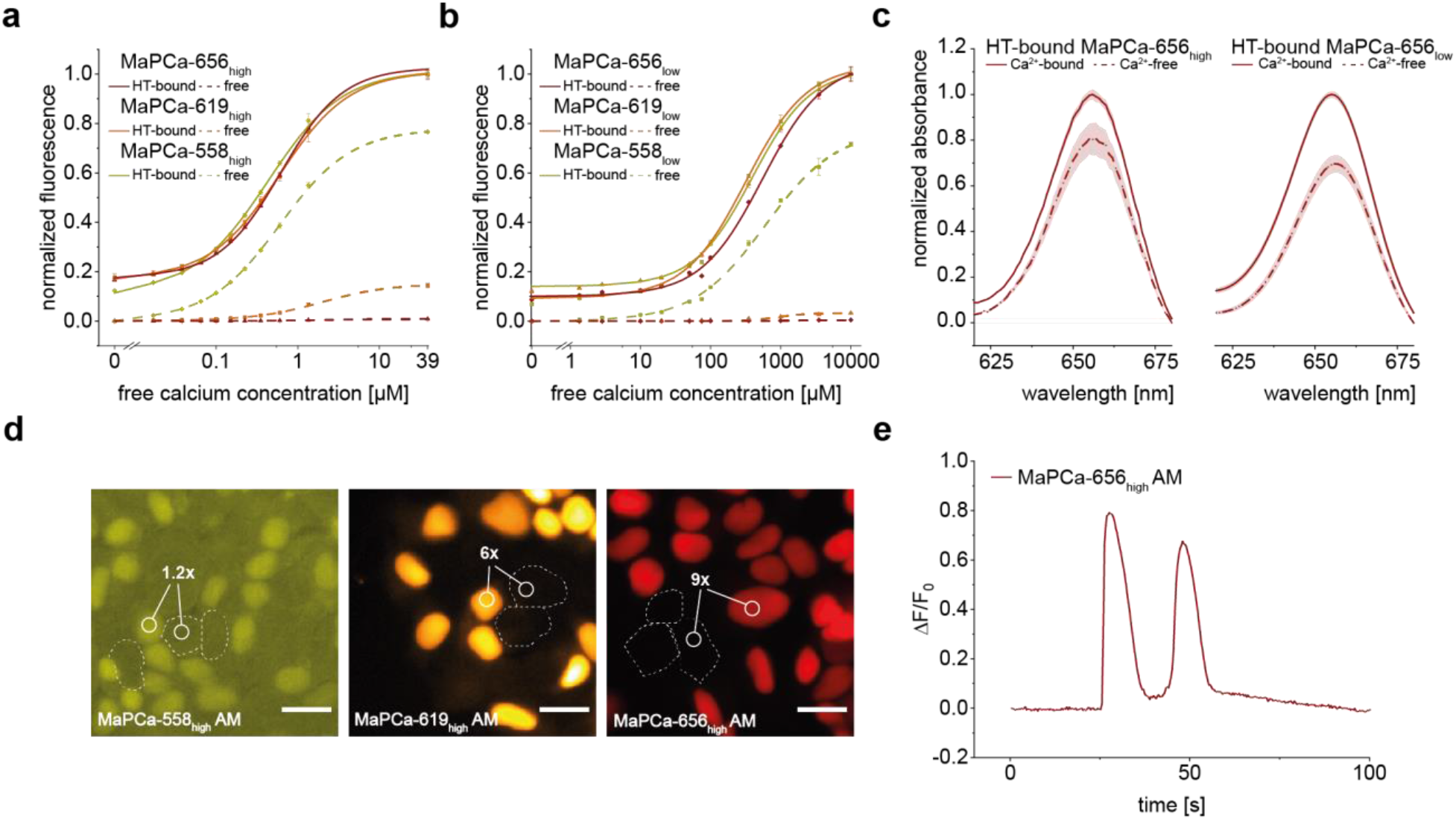
Characterization of MaPCa dyes. (a,b) Calcium titration of (a) MaPCa_high_ and (b) MaPCa_low_. (c) Absorbance spectra of HT-bound MaPCa-656 indicators show calcium dependent absorbance increase. (d) Fluorescence microscopy images of a co-culture of HaloTag-NLS-expressing and non-expressing 293 cells. Cells were incubated with 1 μM MaPCa-558_high_ AM (left), MaPCa-619_high_ AM (middle) or MaPCa-656_high_ AM (right) for 2 h and imaged under no-wash conditions. Turn-on numbers represent average of n = 200 cells. Scale bars: 20 μm. (e) Exemplary fluorescence trace of 293 stably expressing HaloTag-SNAP-tag fusion protein in the nucleus, incubated with MaPCa-656_high_ AM and perfused with 100 μM ATP. HT= HaloTag.

We hypothesized that the increase of fluorescence intensity of the MaPCa dyes upon calcium binding should be mainly due to decreased PET quenching. However, MaPCa-656_high_ and MaPCa-656_low_ show a 20-30% increase in absorbance upon calcium binding (Fig. 3c). This can be rationalized considering that both indicators, when bound to the HaloTag in the absence of calcium, are not fully in the open state. Calcium binding then weakens the electron-donating effect of the aniline moiety, pushing the equilibrium further to the open, fluorescent state (Supporting Fig. 5).

### In-cellulo characterization of the MaPCa dyes

For first cellular calcium imaging experiments, AM-esters of the MaPCa_high_ indicators were applied to co-cultures of 293 cells stably expressing a nuclear localized HaloTag and 293 cells without HaloTag. Imaging the cells without any washing steps after 2 h incubation already revealed efficient HaloTag labeling (Fig. 3d), demonstrating that these molecules are cell permeable. The comparison of the cytosolic background fluorescence intensity in non-expressing cells vs. the nuclear signal of expressing cells reveals that MaPCa-619_high_ AM and MaPCa-656_high_ AM show excellent signal to background ratios (F_nuc_/F_cyt_= 6 and 9, respectively) (Fig. 3d). This can be rationalized by the high fluorogenicity of these two substrates. In contrast, the low fluorogenicity of MaPCa-558_high_ AM results in a high background under no-wash conditions (F_nuc_/F_cyt_= 1.2) (Fig. 3d). Furthermore, all MaPCa_high_ AM indicators translated the calcium concentration rise induced by ATP treatment by a mean fluorescence intensity increase (ΔF/F0) ranging between 0.5 and 2. (Fig. 3e, Supporting Fig. 6).

### MaPCa dyes report on calcium signaling in neurons

In a next step, the performance of the MaPCa indicator series was evaluated in rat primary hippocampal neurons. For experiments with primary neuronal cultures, the possibility to perform the labeling without any washing steps is important, as such steps are known to disturb viability of primary cell cultures.^29^ rAAV transduced rat primary hippocampal neurons expressing HaloTag-mEGFP strictly in the cytoplasm were labeled with either MaPCa-619_high_ AM or MaPCa-656_high_ AM and imaged under no-wash conditions. Both dyes led to efficient and homogeneous HaloTag labeling without the occurrence of a significant background signal or unspecific staining. While usable, MaPCa-558_high_ AM required a washing step to reach results similar to MaPCa-619_high_ AM and MaPCa-656_high_ AM (Fig. 4a, Supporting Fig. 7). To test the sensitivity of the high affinity MaPCa indicators, labeled neurons were stimulated with a distinct number of action potentials (APs) using electric field stimulation.^30^ All dyes allowed the detection of a single AP with ΔF/F0 values ranging between 3% (MaPCa-558_high_ AM) and 6% (MaPCa-656_high_ AM), while ΔF/F0 of 120% was obtained using MaPCa-656_high_ AM with a 160 AP burst (Fig. 4b, Supporting Fig. 8, Supporting Video 1).

**Fig. 4:**
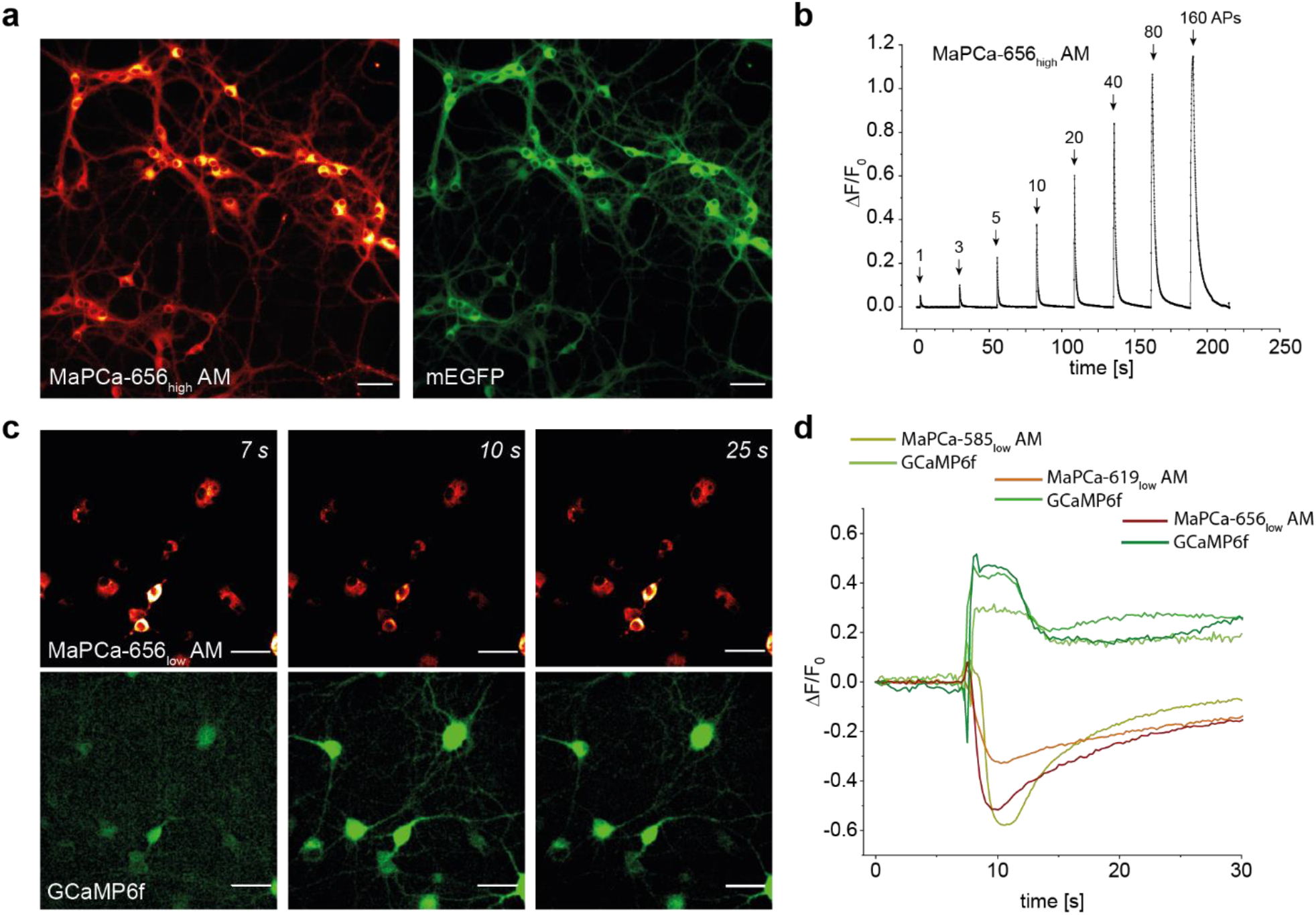
MaPCa dyes can report on calcium flux in primary rat hippocampal neurons. (a) Fluorescence microscopy images of primary rat hippocampal neurons expressing NES-HaloTag-eGFP incubated with 1 μM MaPCa-656_high_ AM and imaged under no-wash conditions; MaPCa-656_high_-channel (left) and eGFP-channel (right). Scale bar 50 μm. (b) Baseline-corrected average trace of stimulated neurons expressing HaloTag and incubated with 1 μM MaPCa-656_high_ AM under no-wash conditions (n ≥ 50 cells). APs: 1,2,5,10,20,40,80,160. (c) Fluorescence microscopy images of rat hippocampal neurons expressing ER-localized HaloTag and cytosolic GCaMP6f. Cells were incubated with 1 μM MaPCa-656_low_ AM for 2 h and imaged under no-wash conditions. After ~7 s caffeine (final conc.: 20 mM) was added. (d) Fluorescence time trace of a representative cell in (c) and of identically treated cells with the indicators MaPCa-558_low_ AM and MaPCa-619_low_ AM (single representative cell) imaged simultaneously with GCaMP6f. Scale bar 50 μm.

The lower calcium affinity of the MaPCa_low_ series allows to report calcium fluctuations in compartments with high basal calcium concentrations such as the ER (Ca^2+^ conc.: ~ 500 μM).^19^ Therefore, the MaPCa dyes were target to the ER through rat hippocampal neuron transduction localizing a HaloTag-SNAP-tag fusion in the ER. Co-staining of SNAP-tag confirmed efficient and specific labeling of HaloTag with MaPCa_low_ dyes under no-wash conditions, with the exception of MaPCa-558_low_ AM that required a washing step to reduce background (Supporting Fig. 9). The ER is a calcium store which, upon stimulation, can release calcium into the cytosol. Here, the RyR2 channel plays a crucial role as a calcium-induced calcium-release channel.^31^ As the red-shifted wavelengths of the MaPCa dyes do not spectrally overlap with the GFP channel, we multiplexed the MaPCa signal from the ER with a cytosolic GCaMP6f, *i.e.* to simultaneously image calcium efflux from the ER and cytosolic influx upon stimulation. Specifically, rat hippocampal neurons were double transduced using rAAVs expressing both constructs individually and then labeled with the MaPCa_low_ AM indicators. Upon addition of caffeine, a RyR2 stimulant,^31,32^ we could simultaneously record a signal decrease in the ER due to calcium efflux (MaPCa_low_ AM) and a concomitant signal increase in the cytosol due to calcium influx (GCaMP6f) (Fig. 4 c,d, Supporting Fig. 10, Supporting Video 2). This demonstrates how MaPCa AM dyes allow, in combination with established GCaMP sensors, to visualize the complex interplay between calcium pools in different cellular compartments in a time-resolved manner.

### Bioluminescence as a readout

The MaPCa dyes could potentially also be used for the labeling of H-Luc, a chimera between HaloTag and the furimazine-dependent luciferase NanoLuc.^33^ Labeling of H-Luc with rhodamine dyes can result in efficient BRET from NanoLuc to the bound rhodamine, such that emission at both 450 nm and at the emission wavelength of the bound rhodamine can be observed. We hypothesized that labeling H-Luc with MaPCa dyes would lead to the development of bioluminescent calcium indicators with tunable emission wavelengths with up to far-red light emission (Fig. 5a). Existing bioluminescent calcium indicators, such as Orange CAMBI,^34^ GLICO,^35^ LUCI-GECO1,^36^ CeNL^37^ or CalfluxVTN^38^ rely exclusively on fluorescent proteins that possess emission maxima restricted below 600 nm. We therefore labeled H-Luc with the MaPCa dyes and recorded the emitted light upon addition of furimazine in the absence and presence of calcium. As is already apparent by eye (Fig. 5b), the color of the emitted light dramatically depends on both, the presence of calcium as well as the nature of the MaPCa dye attached to H-Luc (Fig. 5c). The efficiency of BRET is largest for MaPCa-558_high_ attached to H-Luc, as it has the largest spectral overlap with the BRET donor. As the intensity of the emission of the MaPCa dye depends on the concentration of calcium, measuring the ratio of the intensity of emitted light at 450 nm versus the intensity of the light emitted at the emission maximum of the rhodamine dye can thus be used to record changes in calcium concentrations (Fig. 5b, c, Supporting Fig. 11). The maximal change in ratio ranged from 6.5 for H-Luc labeled with MaPCa-656_high_ to 4.2 for H-Luc labeled with MaPCa-619_high_. The H-Luc-MaPCa ratio changes are comparable to those of previously described ratiometric, bioluminescent calcium sensors and to the best of our knowledge, H-Luc labeled with MaPCa-656_high_ is the first bioluminescent calcium indicator with emission in the far-red. To demonstrate how these ratiometric bioluminescent calcium sensors can be exploited for cellular applications, Flp-In 293 cells with a nuclear H-Luc expression were labeled with the MaPCa_high_ AM dye series. The cells were then exposed to a solution of ATP and thapsigargin and the emission ratio of the emitted light was recorded. A significant change in luminescence emission ratio for all three MaPCa dyes was observed upon drug treatment, the value being the highest for MaPCa-558_high_ AM (1.7-fold) and the smallest for MaPCa-656_high_ AM (1.3-fold) (Fig. 5d). In such given experimental conditions, each channel luminescence intensity was integrated in less than 500 ms, allowing changes in calcium concentrations to be followed with good temporal resolution. The z-factor is a measure for the statistical effect size used to judge the suitability of an assay for high-throughput screening (HTS) approaches. 293-cells expressing H-Luc labeled with MaPCa-558_high_ AM presented a z-value of 0.58 upon ATP/thapsigargin treatment, highlighting the suitability of such bioassay for HTS (z-factors ≥ 0.5 indicate excellent suitability).^39^ Finally, low-affinity bioluminescent calcium indicators could be generated by labeling H-Luc with the MaPCa_low_ indicators, demonstrating the modularity of the approach (Supporting Fig. 12).

**Fig. 5:**
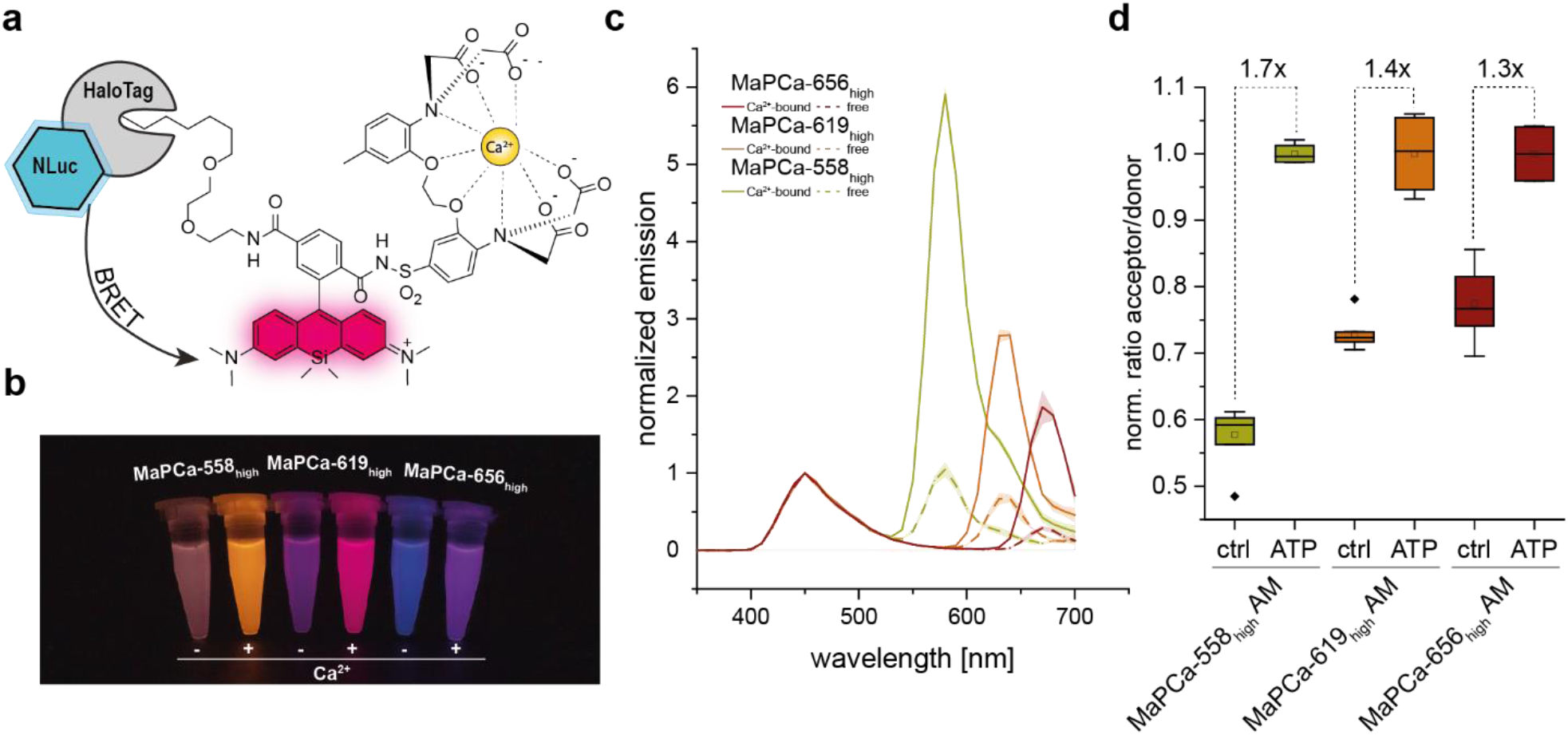
Characterization of MaPCa_high_ based bioluminescent indicators. (a) Bioluminescent H-Luc transfers energy (BRET) to bound MaPCa dyes. (b) Picture of Eppendorf tubes filled with H-Luc labeled MaPCa_high_ dyes in the absence or presence of calcium. (c) Normalized *in vitro* emission spectra of H-Luc labeled MaPCa_high_ dyes, with-and without calcium. (d) Normalized acceptor-donor ratio of 293 cells expressing H-Luc in the nucleus and labeled with 1 μM MaPCa_high_ AM-dyes. Shown is the ratio of control wells and wells treated of 100 μM ATP and 5 μM thapsigargin (N ≥ 4 wells per condition).

## Conclusion

We have introduced a new design principle for the development of localizable and fluorogenic calcium indicators. Using this strategy, we have developed several indicators with different colors, up to the far-red, and with different calcium affinities. What distinguishes these indicators from previous work is the good permeability of the probes and the possibility to use them without additional washing steps to remove unbound indicator. This greatly facilitates their use in most biological applications. Furthermore, they are accessible through a short and modular synthetic pathway. We demonstrated applications of the indicators in rat hippocampal neurons, where the high-affinity indicator MaPCa_high_ could detect single APs under no-wash conditions. The low-affinity indicator MaPCa_low_ was successfully localized in the ER, where it could detect calcium efflux isochronal to increase in cytosolic calcium detected by GCaMP6f. We furthermore developed the first far-red bioluminescent calcium indicator by coupling MaPCa with H-Luc, a bioluminescent HaloTag. The use of H-Luc-MaPCa in cells also demonstrated the possibility to use such bioassays in HTS approaches. These examples underscore the versatility of these calcium indicators and their ease of use.

Finally, the established design principles of these calcium indicators should be transferable to analytes other than calcium.

## Methods

Detailed procedures for the synthesis of all compounds, their characterization and imaging experiments are given in the Supplementary Information.

### General

For all microscopy images, a DMi8 widefield microscope (Leica) equipped with a HC PL APO 20x/0.8 (dry) was used. Excitation/Emission settings: TMR-based dyes: λex = 515/30 nm, detection λdet = 609/54 nm; CPY/SiR-based dyes: λex = 635/18 nm, detection λdet = 700/75 nm. Imaging data was processed using Fiji.40 ROIs were drawn manually and average fluorescence intensities extracted and plotted using Origin software. HaloTag in this work corresponds to the variant HaloTag7.

### Signal/background measurements Flp-In 293 cells

Flp-In 293 cells stably expressing HaloTag-SNAP-NLS (see Supporting Information) were co-cultured with non-expressing cells on a 10-well-plate (Greiner bio-one; cellview cell culture slide; glass bottom). After 2 h incubation with 1 μM MaPCa_high_ AM and 0.04% Pluronic-F127, cells were imaged without any washing steps. For analysis, n ≥ 200 cells from at least 4 different wells were selected. Raw intensities over ROI of identical size were averaged (AI) for expressing (AI_E_) and non-expressing (AI_NE_) cells and signal/background were calculated by measuring the ratio AI_E_/AI_NE_.

### ATP perfusion experiments

Flp-In 293 stably expressing HaloTag-SNAP-NLS (see Supporting Information) were seeded in Ibidi μ-Slides (VI 0.4, Poly-L-Lysine coated, part No: 80604) 24 h before start of the experiment. SNAP-tag was co-stained by incubating cells with TMR-CP or SiR-BG at 1 μM for 30 min to 1h at 37°C. After washing twice with medium, cells were stained with calcium dyes (1 μM + 0.04% Pluronic-F127, 2 h) through HaloTag labeling. The chamber was placed on the microscope and attached to a gravity-flow perfusion system. Cells were perfused with HBSS and, upon trigger, with HBSS containing 100 μM ATP. Image acquisition was performed every 350 ms.

### Live-cell labeling and imaging in primary hippocampal neurons

Primary hippocampal neurons were prepared from postnatal P0-P2 Wistar rats as previously described41 (see also supplementary information) and cultured in 24-well glass-bottom plates for 14-18 days. The procedure was conducted in accordance with the Animal Welfare Law of the Federal Republic of Germany (Tierschutzgesetz der Bundesrepublik Deutschland, TierSchG) and the regulation for animals used in experiments (1 August 2013, Tierschutzversuchsverordnung). To euthanize rodents for the subsequent preparation of any tissue, all the regulations given in Section 4 of the TierSchG were followed. As the euthanization of animals is not an experiment on animals according to Section 7 paragraph 2 sentence 3 of the TierSchG, no specific authorization or notification is required. On day 7-8, neurons were transduced with 0.5 μL of the corresponding rAAVs for hSyn1 driven expression of NES-HaloTag-mEGFP; CalR-HaloTag-Snap-tag-KDEL (ER-localizing) or GCaMP6f (Addgene 100837-AAV1). On day 14-18 the neurons were incubated for at least 2 h with 1 μM of the corresponding MaPCa dyes containing 0.04% Pluronic-F127. While the medium of MaPCa-558 was subsequently exchanged once, the other dyes were imaged under no-wash conditions. In case a co-stain was utilized, this was added before the MaPCa-staining at 1 μM for 1 h followed by a single medium exchange afterwards. For stimulation of the ER-localized MaPCa dyes, a caffeine-solution (final concentration: 20 mM) was administered manually during recording (Frame rate: 250 ms). For electric field stimulation, synaptic blockers NBQX (10 μM, Santa Cruz) and APV (25 μM, Sigma Aldrich) were added in order to suppress natural spiking activity. Then, a custom-built30 electrode was inserted into the wells and APs (1,2,5,10,20,40,80,160: with 25 s pauses) were evoked with following settings: Pulse width: 1 ms; Amperage 100 mA; Frequency 80 Hz. Data represents the average of N ≥ 50 manually selected ROI of different neurons from at least three different wells. Frame rate: 50 ms.

### Live-cell measurements with bioluminescent readout

The Flp-In System (ThermoFisher Scientific) was used to generate stable 293 cells expressing H-Luc in the nucleus (see Supporting Information). The cells stably expressing H-Luc-NLS were plated on a black flat glass bottom 96-well plates (Eppendorf; tissue culture treated). They were incubated with 1 μM MaPCa_high_ AM dyes and 0.04% Pluronic-F127 for at least 2 h in 100 μL at 37°C. 50 μL of a mix substrate/ extracellular inhibitor were added in each well to measure only the luminescence coming from intact cells [NanoBRET™ Nano-Glo® Substrate/ Extracellular NanoLuc® Inhibitor solution (final dilution substrate: 1’000x; final dilution inhibitor: 3’000x)]. Half the wells were treated with an additional 10 μL of mixture ATP/thapsigargin (final concentrations ATP: 100 μM; thapsigargin: 5 μM). Measurements were performed stepwise (max three wells at a time, max two minutes delay between pipetting and imaging) in order to ensure identical conditions. Spectra from whole sample wells were measured on a plate reader (Spark20M, Tecan). Data represents averaged results from N=4 experimental replicates with standard deviations. The average ratios between BRET acceptor and BRET donor of ATP/thapsigargin-treated wells were then compared to the non-treated wells.

The z-factor was calculated with the formula

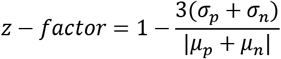

With σ = standard deviation; μ = mean; p = positive control and n = negative control.

### General statistics

All measurements were measured at least in triplicates and presented as mean with s.d. if not stated otherwise. Cellular imaging experiments were performed at least twice on different days.

## Supporting information

Supplemental Imformation

## Acknowledgements

N.M and M.-C.H. are grateful for a Boehringer Ingelheim Fonds PhD Fellowship. This work was furthermore supported by the Max Planck Society and the Heidelberg Biosciences International Graduate School (HBIGS). N.M. was further supported by SFB grant 1129. The authors furthermore want to thank Bettina Réssy and Dominik Schmidt for the synthesis of starting materials, Andrea Bergner for plasmid/protein productions and purifications and Annette Herold for the rAAV production. Furthermore, the authors want to thank Elisa D’Este, Jasmine Hubrich, Angel Rafael Cereceda Delgado and Victor Macarrón Palacios for the support in neuronal cell culture.

## Contributions

N.M. and K.J. planned the experiments and co-wrote the paper with the help of all authors. N.M. synthesized and characterized the MaPCa_high_ dyes and performed all imaging experiments. M.B. synthesized and characterized MaPCa_low_. M.-C.H. and C.-M.G. assisted with neuronal cell culture experiments. C.H. helped to develop the synthetic pathway. S.K. and B.K generated cell lines. J.H. contributed to planning and execution of the bioluminescence experiments.

## Additional Information

Correspondences and requests for material should be addressed to K.J.

