## Supplemental Imformation for "Fluorescent and bioluminescent calcium indicators with tuneable colors and affinities"

### Supplementary Figures and Tables

| | | |
|---|---|---|
| 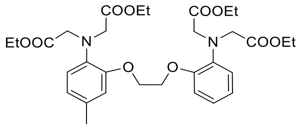 | 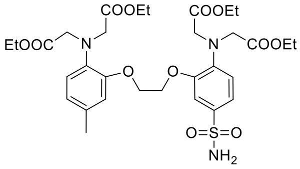 | 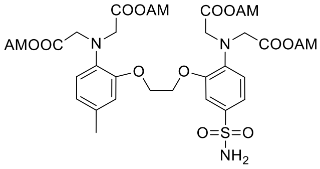 |
| **01** | **03** | **05** |
| 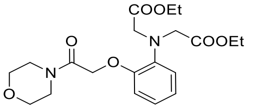 | 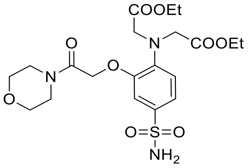 | 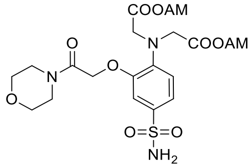 |
| **02** | **04** | **06** |
| 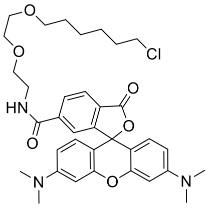 | 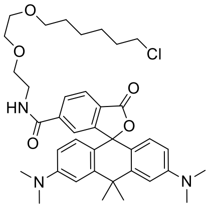 | 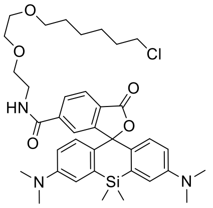 |
| **TMR-CA** | **CPY-CA** | **SiR-CA** |
| 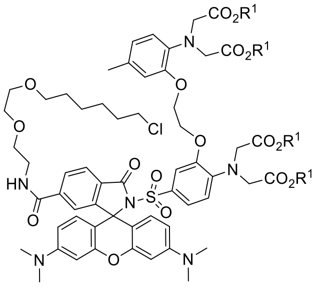 | 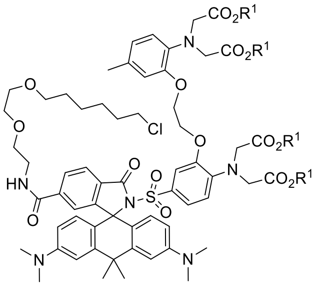 | 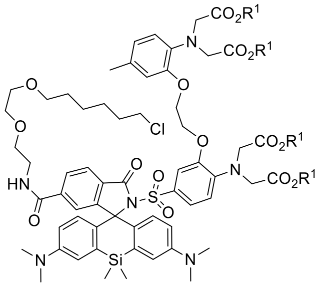 |
| $R^1$= H **\| MaPCa-558$_{high}$**<br>$R^1$= AM **\| MaPCa-558$_{high}$ AM** | $R^1$= H **\| MaPCa-619$_{high}$**<br>$R^1$= AM **\| MaPCa-656$_{high}$ AM** | $R^1$= H **\| MaPCa-656$_{high}$**<br>$R^1$= AM **\| MaPCa-656$_{high}$ AM** |
| 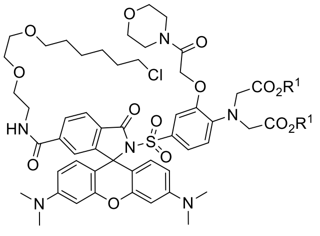 | 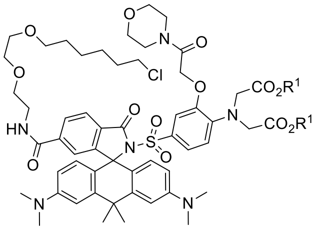 | 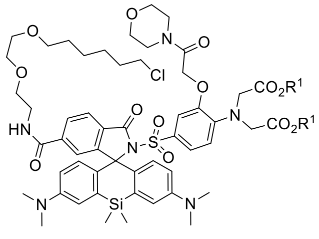 |
| $R^1$= H **\| MaPCa-558$_{low}$**<br>$R^1$= AM **\| MaPCa-558$_{low}$ AM** | $R^1$= H **\| MaPCa-619$_{low}$**<br>$R^1$= AM **\| MaPCa-619$_{low}$ AM** | $R^1$= H **\| MaPCa-656$_{low}$**<br>$R^1$= AM **\| MaPCa-656$_{low}$ AM** |

**Supporting Fig. 1:** Structure of main synthetic molecules in this work.

Synthesis of BAPTA-Sulfonamide:

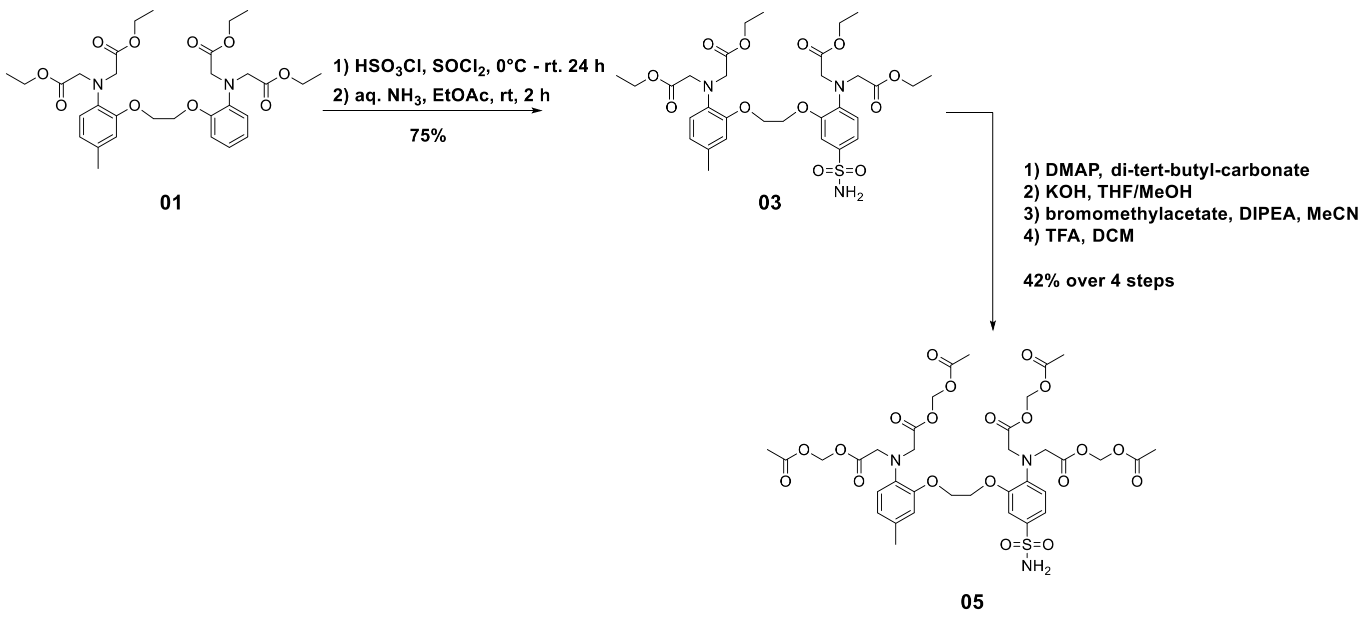

Synthesis of MOBHA-Sulfonamide:

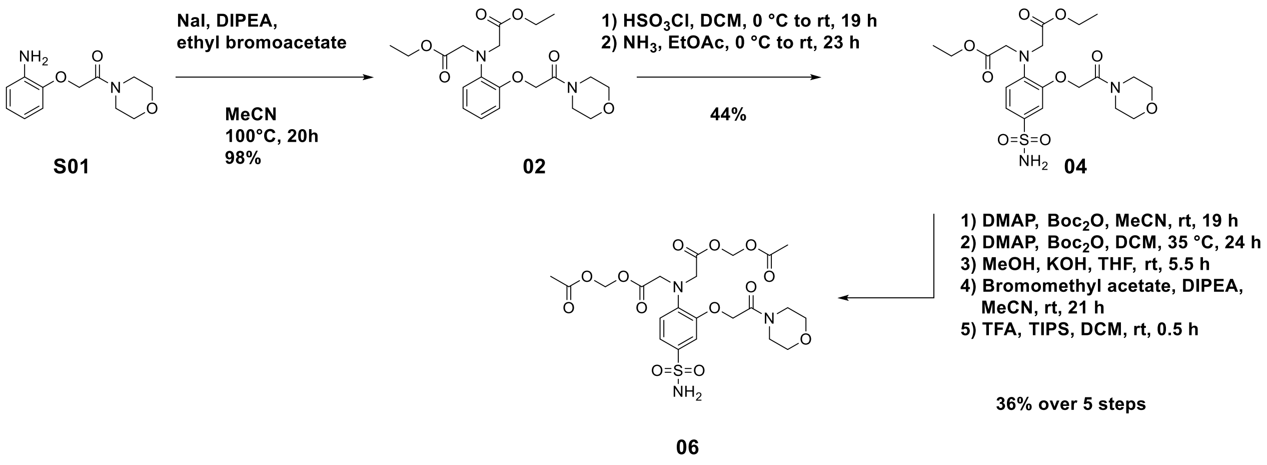

**Supporting Fig. 2:** Synthetic pathways to BAPTA- and MOBHA-sulfonamide.

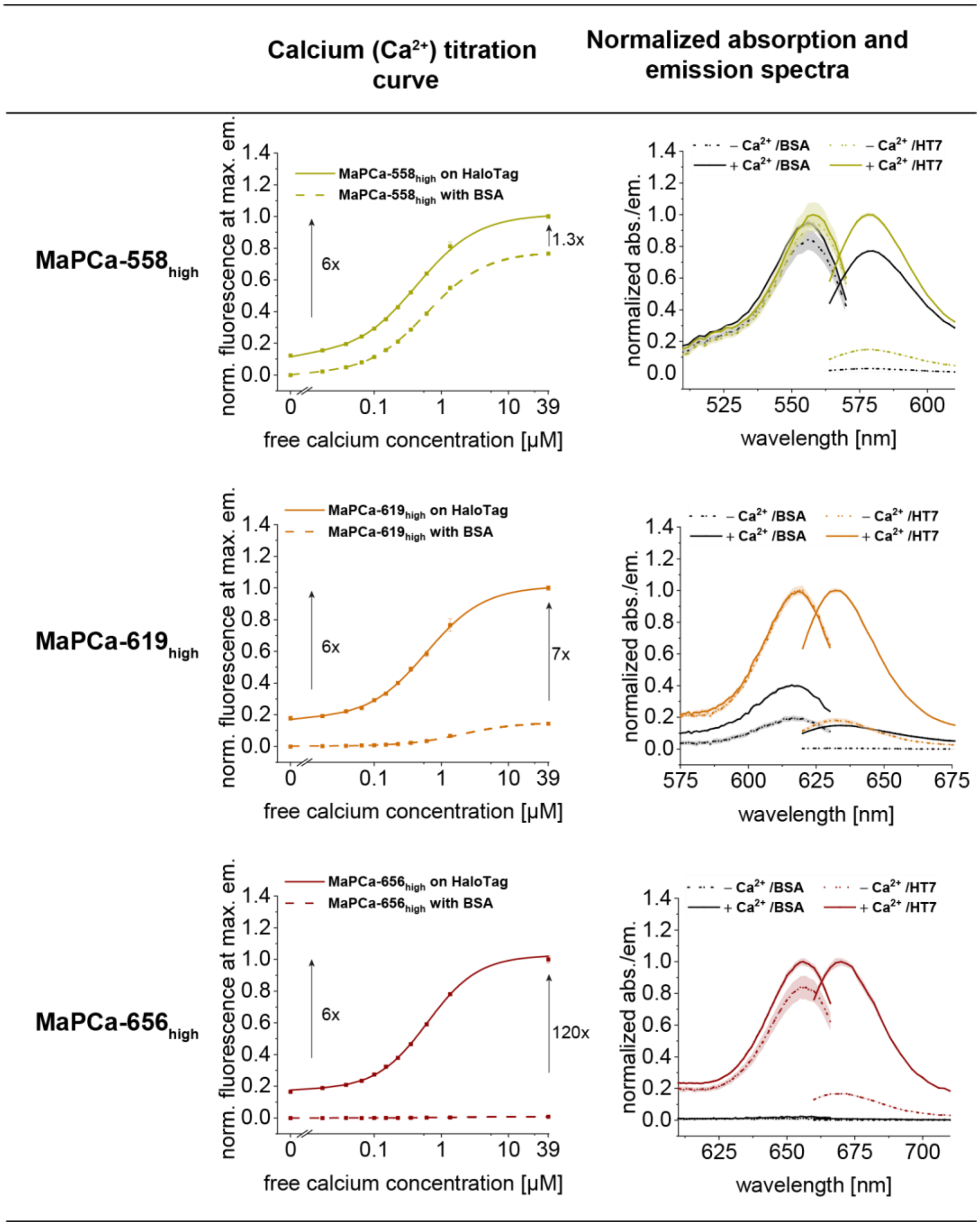

**Supporting Fig. 3:** Calcium titration curves and absorption/emission spectra for $MaPCa_{high}$ dyes. $K_D(Ca^{2+})$-values are given in Supporting Table 1.

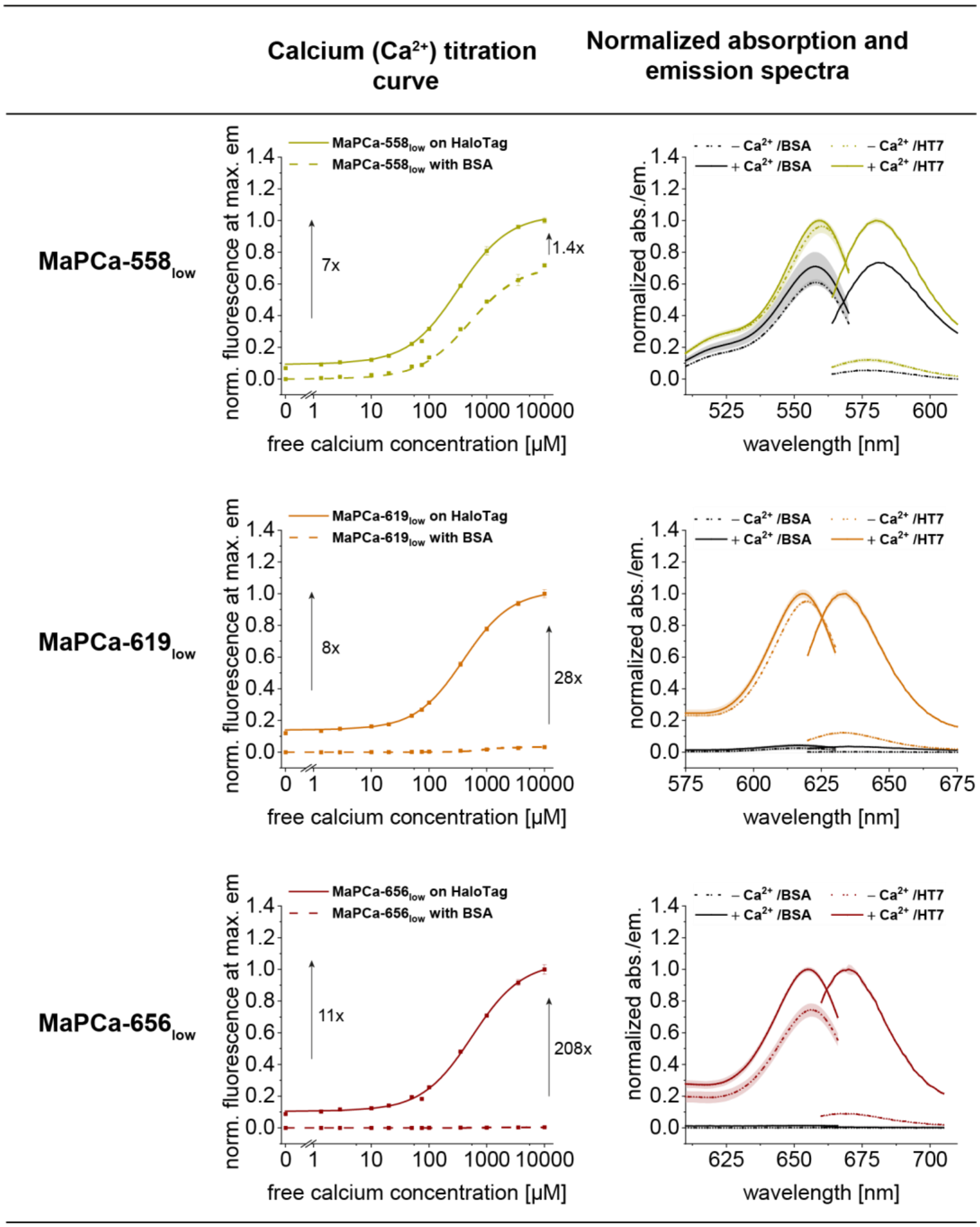

**Supporting Fig. 3:** Calcium titration curves and absorption/emission spectra for $MaPCa_{low}$ dyes. $K_D(Ca^{2+})$-values are given in Supporting Table 1.

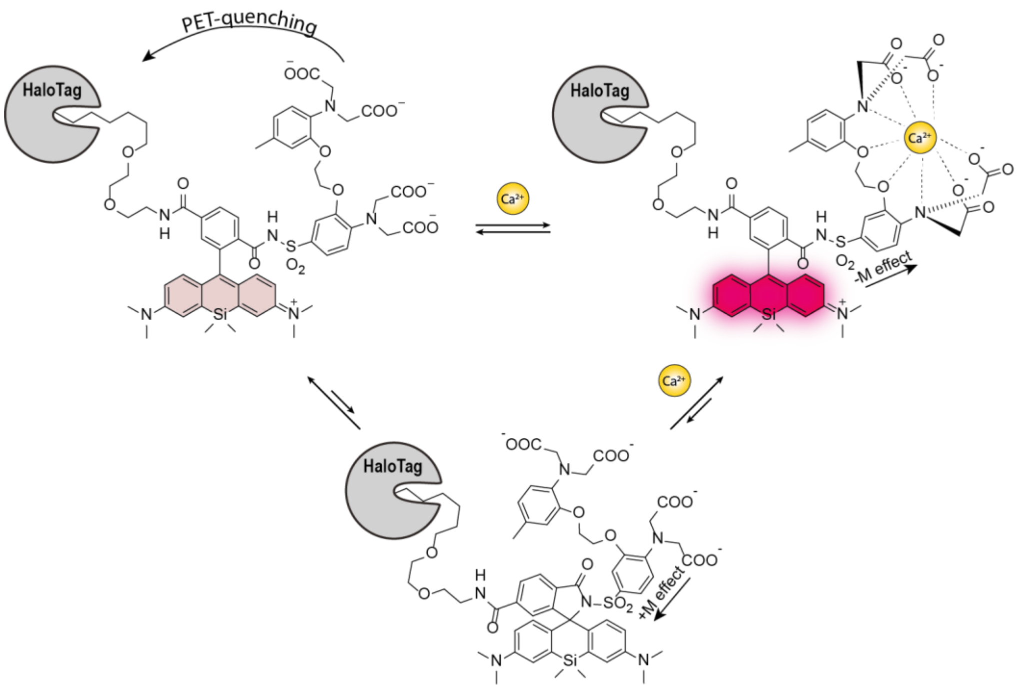

**Supporting Fig. 4:** Schematic representation of the quenching mechanism *via* the influence of calcium on the open/close equilibrium (example of MaPCa-656$_{high}$). The two upper structures do absorb light at 656 nm, while the lower, cyclized structure does not.

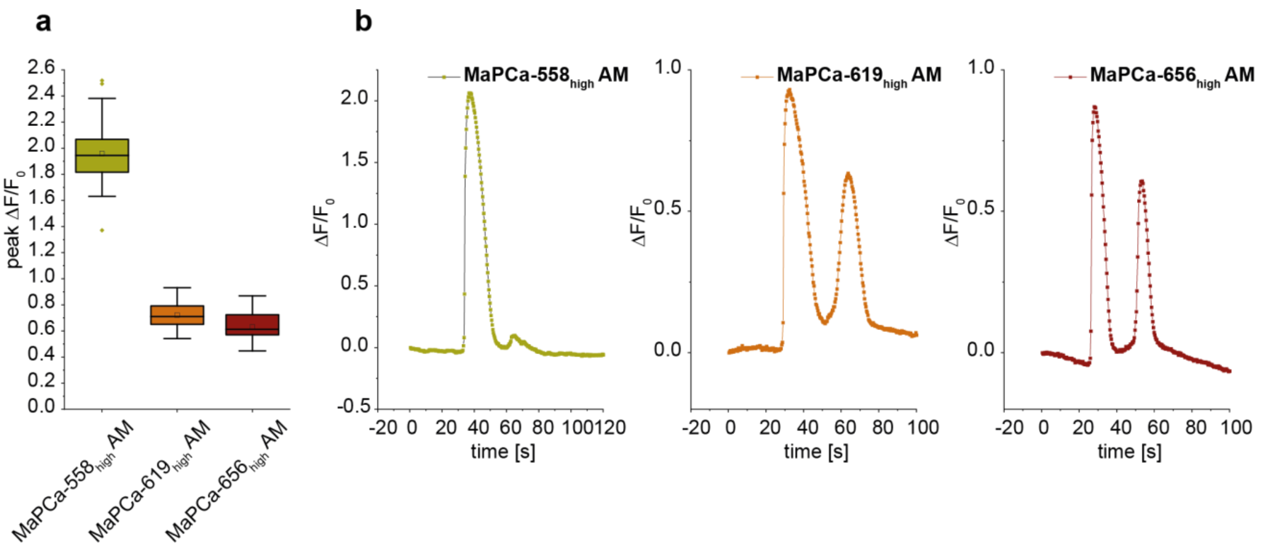

**Supporting Fig. 5:** Analysis of microscopy data of Flp-In 293 cells stably expressing a Halo-SNAP-NLS construct. Live cells were perfused with an HBSS solution and stimulated by fluid exchange with HBSS containing 100 µM ATP. (a) peak $\Delta F/F_0$ of the different MaPCa$_{high}$ AM-indicators upon ATP perfusion (n ≥ 50 cells). (b) Exemplary traces of the ATP-perfusion of MaPCa-558$_{high}$ AM (left), MaPCa-619$_{high}$ AM (middle) and MaPCa-656$_{high}$ AM (right). Frame rate: 350 ms.

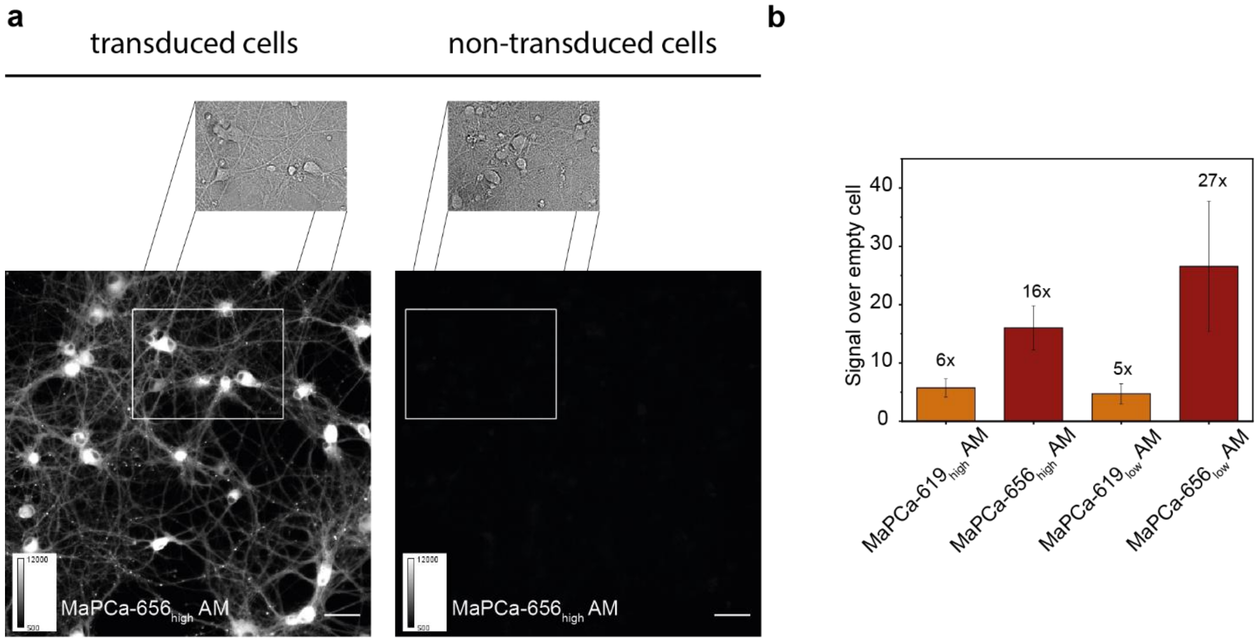

**Supporting Fig. 6:** The MaPCa dyes enable no-wash labeling of cytosolic HaloTag expressed in primary rat hippocampal neurons. (a) Widefield fluorescence microscopy data from primary rat hippocampal neurons expressing cytosolically NES-HaloTag-mEGFP and non-transduced cells. Example no-wash fluorescence with brightfield-inset of transduced (left) and non-transduced (right) neurons incubated for 2 h with 1 µM MaPCa-656$_{high}$ AM. Scale bars 50 µm. (b) Histogram representing the signal over empty cell signal of the different MaPCa-dyes. N ≥ 30 cells.

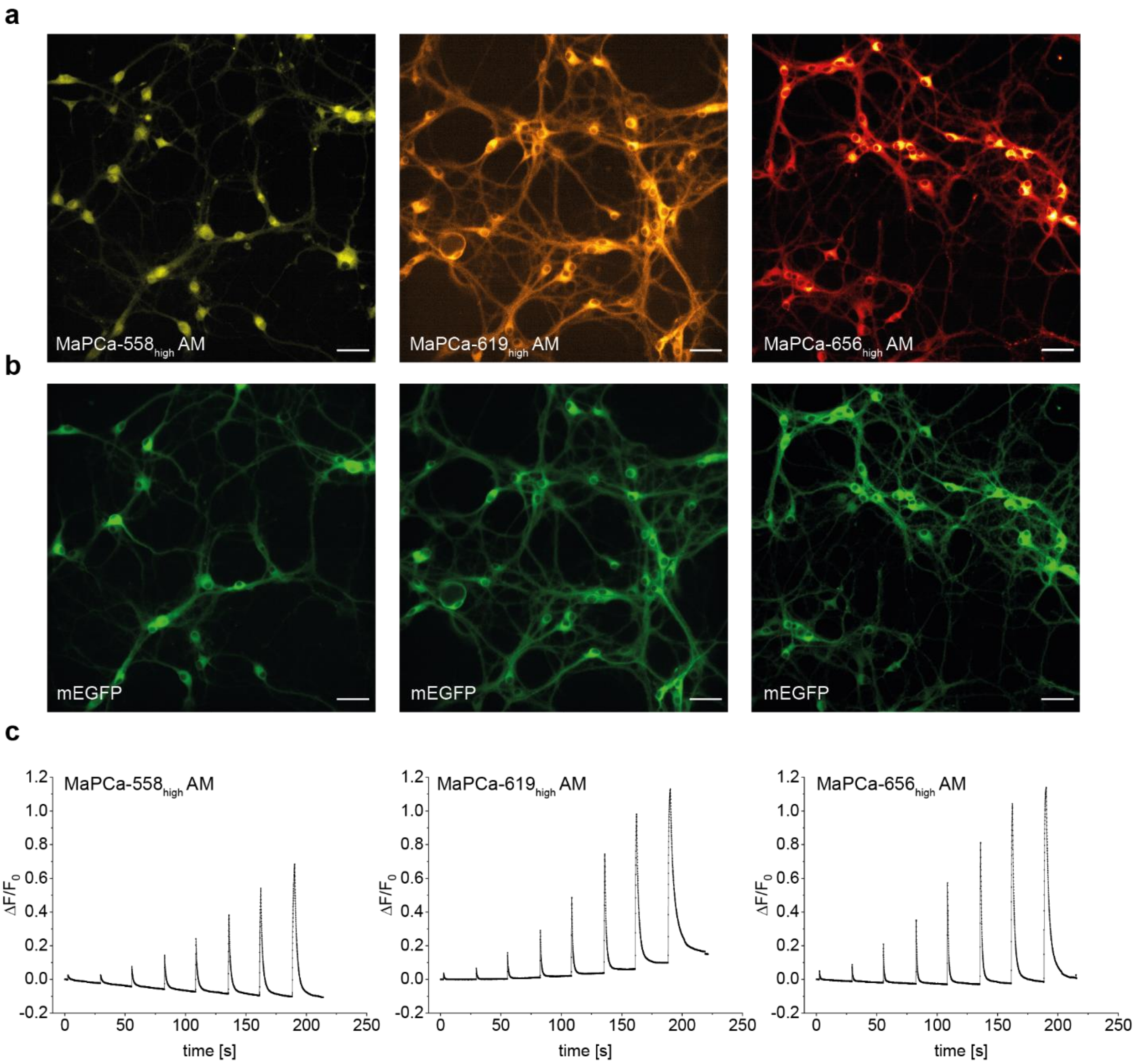

**Supporting Fig. 7:** MaPCa dyes reliably detect induced action potentials in primary rat hippocampal neurons. Widefield microscopy images from primary rat hippocampal neurons (DIV 14-18), expressing NES-HaloTag-mEGFP in the cytosol, incubated with 1 µM MaPCa$_{high}$ AM for 2 h. (a) Microscopy images of MaPCa-558$_{high}$ AM (left, washed once), MaPCa-619$_{high}$ AM (middle, no-wash) and MaPCa-656$_{high}$ AM (right, no-wash). (b) GFP-channel of (a). (c) Averaged non-corrected traces of electric field stimulation of MaPCa-558$_{high}$ AM (left, washed once), MaPCa-619$_{high}$ AM (middle, no-wash) and MaPCa-656$_{high}$ AM (right, no-wash). N ≥ 50 cells. APs: 1,2,5,10,20,40,80,160. Frame rate: 50 ms. Scale bars: 50 µm.

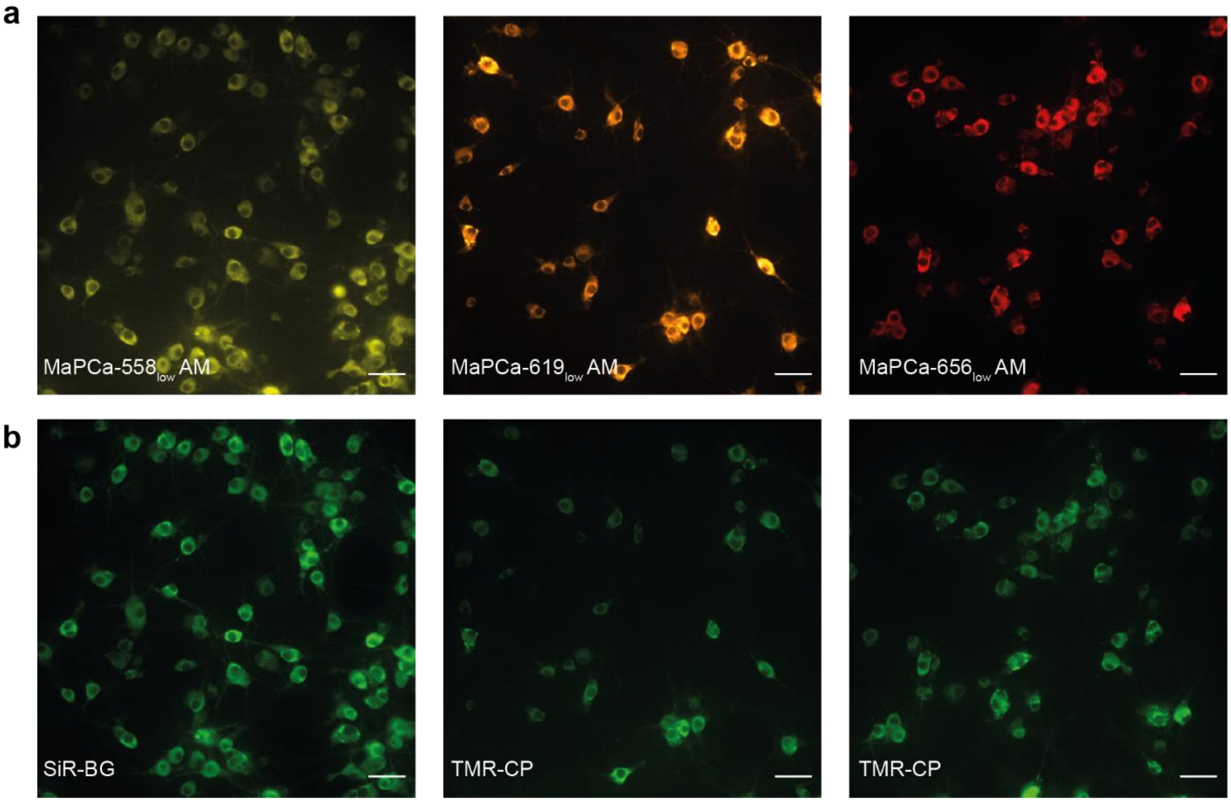

**Supporting Fig. 8:** MaPCa$_{low}$ bound to ER-localized HaloTag. Widefield microscopy data from primary rat hippocampal neurons expressing a HaloTag-SNAP construct in the ER, incubated with 1 µM of counterstain for 1 h, washed, then incubated with MaPCa$_{low}$ AM for 2 h at 1 µM. (a) Microscopy images of MaPCa-558$_{low}$ AM (left, washed once), MaPCa-619$_{low}$ AM (middle, no-wash) and MaPCa-656$_{low}$ AM (right, no-wash). (b) counterstain images with SNAP-tag substrates. Left: SiR-BG; middle: TMR-CP; right: TMR-CP. Scale bars 50 µm.

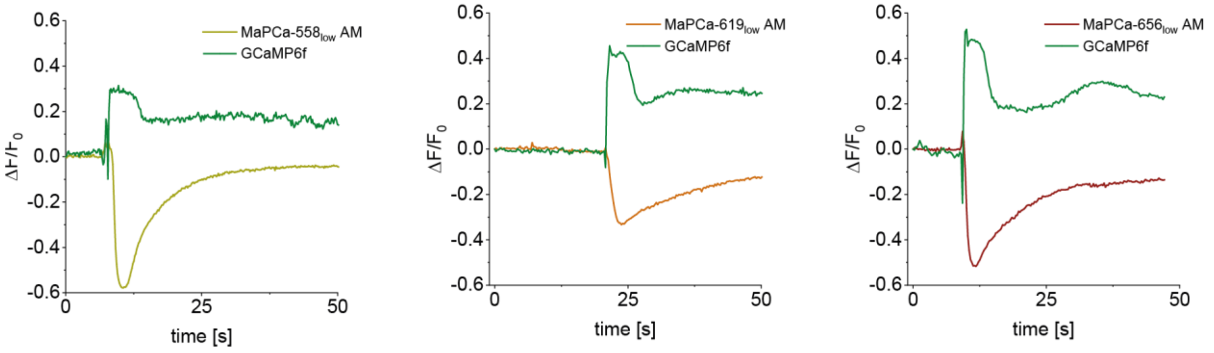

**Supporting Fig. 9**: Representative single-cell fluorescence time trace of rat hippocampal neurons expressing ER-localized HaloTag7 and cytosolic GCaMP6f. Cells were incubated with 1 µM $MaPCa_{low}$ AM-dyes for 2 h and imaged under no-wash conditions or washed once (MaPCa-$558_{low}$ AM). After several seconds caffeine (final conc.: 20 mM) was added.

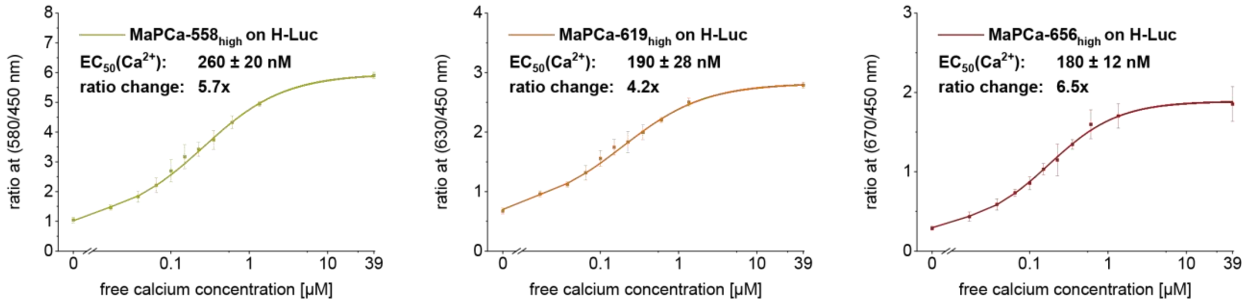

**Supporting Fig. 10:** $MaPCa_{high}$ dyes in combination with H-Luc detect calcium changes. *In vitro* calcium titrations of H-Luc labeled with $MaPCa_{high}$ indicators. Plotted are the ratios of BRET acceptor/donor.

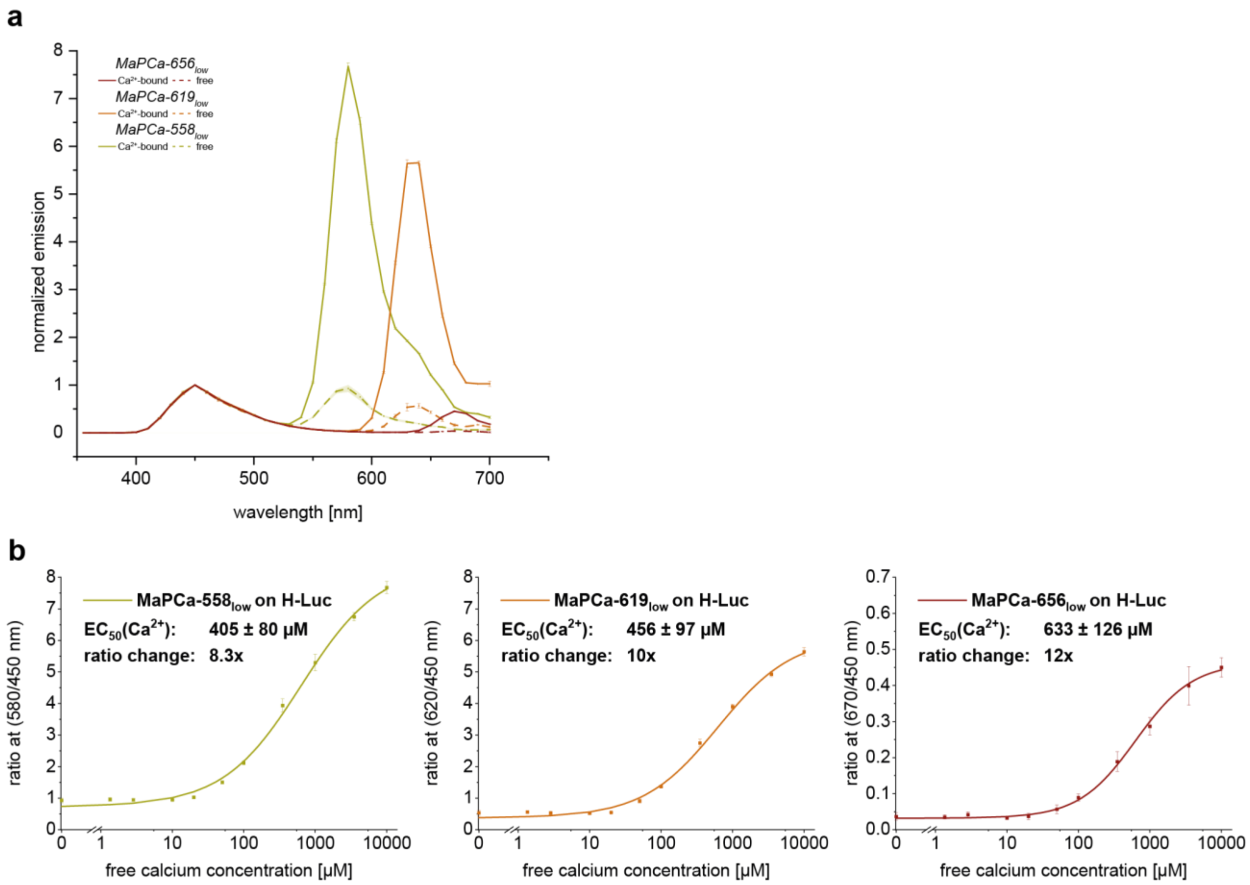

**Supporting Fig. 11:** MaPCa$_{low}$ dyes in combination with H-Luc *in vitro.* (a) Normalized *in vitro* emission spectra of H-Luc labeled MaPCa$_{low}$ dyes, with-and without calcium. (b) *In vitro* calcium titrations of H-Luc labeled with MaPCa$_{low}$ indicators. Plotted are the ratios of BRET acceptor/donor.

**Supporting Table 1:** Spectroscopic data of MaPCa-dyes.

| | $F/F_0$ | | | | $\lambda$Ex/$\lambda$Em [nm] | $K_D(Ca^{2+})$ [μM]* | | ε ($mM^{-1}cm^{-1}$) | | | | Φ | | | |
|---|---|---|---|---|---|---|---|---|---|---|---|---|---|---|---|
| | $Ca^{2+}$-bound | $Ca^{2+}$-free | HT-bound | HT-free | | | | | | | | | | | |
| | HT-binding | HT-binding | $Ca^{2+}$-binding | $Ca^{2+}$-binding | | HT | BSA | HT | | BSA | | HT | | BSA | |
| | | | | | | | | $+Ca^{2+}$ | $-Ca^{2+}$ | $+Ca^{2+}$ | $-Ca^{2+}$ | $+Ca^{2+}$ | $-Ca^{2+}$ | $+Ca^{2+}$ | $-Ca^{2+}$ |
| MaPCa-$558_{high}$ | 1.3 | 4 | 6 | 22 | 558/580 | 0.41 | 0.57 | 81 | 79 | 71 | 68 | 49% | 7% | 30% | 2% |
| MaPCa-$619_{high}$ | 7 | 30 | 6 | 24 | 619/632 | 0.57 | 2.20 | 91 | 91 | 34 | 18 | 60% | 11% | 59% | 4% |
| MaPCa-$656_{high}$ | 120 | 615 | 6 | 31 | 656/670 | 0.58 | 1.80 | 81 | 75 | <2 | <2 | 41% | 7% | NA | NA |
| MaPCa-$558_{low}$ | 1.4 | 2 | 7 | 8 | 560/580 | 224 | 487 | 68 | 66 | 55 | 45 | 38% | 6% | 30% | 3% |
| MaPCa-$619_{low}$ | 28 | 33 | 8 | 10 | 618/633 | 322 | 806 | 93 | 83 | 5 | 3 | 48% | 5% | 40% | 4% |
| MaPCa-$656_{low}$ | 208 | 153 | 11 | 8 | 655/670 | 457 | 926 | 34 | 24 | <2 | <2 | 44% | 5% | NA | NA |

* $K_D(Ca^{2+})$-values were calculated using linear regression of log $[(F - F_{min})/(F_{max} - F)]$ vs. log $[Ca^{2+}]$, where x-intercept= log $K_D(Ca^{2+})$.

**Supporting Table 2:** Plasmids and stable cell lines generated for this work.

| Addgene# | Plasmid | Gene | Stable cell lines |
|---|---|---|---|
| | pcDNA5 | HaloTag7-SNAP-NLS | Flp-In 293 TREx |
| | pcDNA5 | H-Luc-NLS | Flp-In 293 TREx |
| | pAAV2-hSyn | AAV2/1-hSyn-NES-HT7-mEGFP-WPRE-SV40 | - |
| | pAAV2-hSyn | AAV2/1-hSyn-CalR-HT7-Pro30-SNAP-KDEL-WPRE-SV40 | - |
| 100837-AAV1 | | AAV1-Syn-GCaMP6f-WPRE-SV40 | - |

**Supporting Table 3:** Titers of applied rAAVs.

| rAAV(1)-plasmid | titer |
|---|---|
| pGP-AAV2-hSyn2-GCaMP6f-WPRE | 2.8x10^13 GC/mL |
| NES-HaloTag-mEGFP | 3.95x10^13 GC/mL |
| CalR-HaloTag-SNAP-KDEL | 3.79x10^13 GC/mL |

### General Experimental Information

Common reagents were purchased from commercial suppliers (Acros, Bachchem, Fluka, Fluorochem, Merck, Roth, Sigma-Aldrich, TCI and Enamine) and used without further purification. BAPTA-Me **(01)**, SiR-CA, CPY-CA and TMR-CA were synthesized according to literature procedures[1-4] by N. Mertes, B. Réssy or D. Schmidt. The composition of mixed solvents are given by the volume ratio (v/v). Reactions in the absence of air were performed in oven-dried glassware under argon atmosphere.

Reaction progress was either monitored by thin layer chromatography (TLC) on TLC-aluminium sheets (Silica gel 60 $F_{254}$, Merck) or liquid chromatography-mass spectrometry (LCMS- Shimadzu MS2020 connected to a Nexera UHPLC system equipped with a C18 80 Å 1.9 µm, 2.1x50 mm column by Supelco). A typical gradient was from 10% to 90% B within 6 min using a 1 mL/min flowrate (solvent A: 0.1% formic acid in water; solvent B: 0.1% formic acid in acetonitrile). Flash column chromatography was performed on an automated system (Biotage Isolera One) using pre-packed columns (12-40g, 130-400 mesh, Silicycle). Preparative RP-HPLC was performed on an UltiMate 3000 system (Thermo Fisher Scientific) or Waters e2695 system using either a C18 5 µm, 10x250 mm column (flow rate 4 mL/min, Supelco) or a C18 5 µm, 21.2 x 250 mm column (flow rate 8 mL/min, Supelco). A typical run was over 60 min, solvent ratios are given in the procedures (solvent A: 0.1% TFA in water, solvent B: MeCN). For large batches, a preparative Shimadzu-system equipped with an SPD-M20A diode array detector for product visualization and an LCMS-2020 for mass detection on either a Shimadzu Shim-pack GIS C18 column (5 µm, 30 x 250 mm) or on a Shimadzu Shim-pack GIS C18 column (5 µm, 50 x 250 mm) was used. Solvent A: 0.1% FA in water, solvent B: 0.1% FA in MeCN. Compounds were dried under high vacuum using a lyophilizer (Christ) equipped with a vacuum pump (Vacuubrand).

$^{1}H$ and $^{13}C$ NMR spectra were recorded on a Bruker Ascend™ 400 at 400 MHz ($^{1}H$) or 101 ($^{13}C$) MHz. All spectra were recorded at 298 K. Chemical shifts δ are reported in ppm downfield from tetramethylsilane using the residual deuterated solvent signals as an internal reference ($CDCl_3$: δH = 7.26 ppm, δC = 77.16 ppm; DMSO-d6: δH = 2.50 ppm, δC = 39.52 ppm; $CD_3CN$: δH = 1.94 ppm, δC = 118.26 ppm). Chemical shifts δ are given in ppm, coupling constants J in Hertz (Hz). Multiplicities are abbreviated as follows: s = singlet, d = doublet, t = triplet, q = quartet, quin = quintet, m = multiplet and bs = broad signal. Data was processed using Mnova from Mestrelab.

High-resolution mass spectrometry was performed by the MS-facility of the Max Planck Institute for Medical Research on a Bruker maXis IITM ETD mass spectrometer coupled to a Shimadzu Nexera system controlled via o-TOF-Control 4.1 and Hystar 4.1 SR2 (4.1.31.1) software (Bruker). The acquisition rate was set to 3 Hz. The following source parameters were used for positive mode electrospray ionization (ESI+): End plate offset = 500 V; capillary voltage = 3800 V; nebulizer gas pressure = 45 psi; dry gas flow = 10 L/min; dry temperature = 250°C. As transfer, quadrupole and collision cell settings are mass range dependent, they were fine-tuned with consideration of the respective analyte's molecular weight. For internal calibration sodium format clusters were used. Samples were desalted via fast liquid chromatography (TitanTM C18 UHPLC Column (Supelco), 1.9 µm, 80 Å pore size, 20 × 2.1 mm, 2 min gradient from 10 to 98% aq. MeCN with 0.1% FA). Sample dilution in 10% aq. MeCN and injection volumes were chosen depending on the analyte's ionization efficiency (on-column loadings: 0.25–5.0 ng). Automated internal re-calibration and data analysis of the recorded spectra were performed with DataAnalysis 4.4 SR1 software (Bruker).

#### Cloning, Protein Expression and Purification

**General.** Plasmids encoding HaloTag-SNAP for bacterial and mammalian cell protein production at different sub-cellular localizations of mammalian cells (NLS, cyto, ER) were previously described.[5,6] The gene encoding H-Luc was cloned by Gibson assembly[7] in the pCDNA5-FRT vector for mammalian expression (ThermoFisher Scientific) with and without a nuclear localization sequence (NLS, nucleus). For rAVV preparation, mEGFP-HaloTag, SNAP-Halo and H-Luc were analogously cloned in plasmid pAAV2-hSyn paying particular attention to ITR integrity verified by Sanger sequencing (Eurofins) after MaxiPrep DNA extraction (Qiagen). H-Luc was cloned in a pET51b(+) vector (Novagen) featuring an N-terminal His10 for protein purification.

**Protein production and purification.** HaloTag7 and H-Luc proteins were produced and purified as previously explained.[8] In short, the proteins were expressed in the *Escherichia coli* strain BL21(DE3)-pLysS (Novagen) in lysogeny broth (LB)[9] cultures grown at 37°C to an optical density at 600 nm (OD600nm) of 0.8, induced by addition of 0.5 mM isopropyl-β-D-thiogalactopyranoside (IPTG) and grown at 17°C overnight in the presence of 1 mM $MgCl_2$. Cells were harvested by centrifugation (4,500 g, 10 min, 4ºC), lysed by sonication and the lysate was cleared by centrifugation (75,000g, 10 min, 4°C). The proteins were purified using HisPur Ni-NTA Superflow Agarose (Thermo Fisher Scientific, Waltham, MA, USA) by batch incubation followed by washing and elution steps on a polypropylene column (Qiagen). The proteins were subsequently buffer exchanged using a HiPrep 26/10 Desalting column (Cytiva) on an ÄktaPure FPLC to HEPES 50 mM, NaCl 50 mM pH 7.3 (*i.e*. activity buffer). Proteins were concentrated using Ultra-15 mL centrifugal filter devices (Amicon, Merck KGaA, Darmstadt, Germany) with a molecular weight cut-off (MWCO) smaller than the protein size to a final concentration of 500 µM. Proteins were aliquoted and stored at -80°C after flash freezing in liquid nitrogen or kept in glycerol 45% (w/v) at -20°C. Correct size and purity of proteins were assessed by SDS-PAGE and liquid chromatography-mass spectrometry (LC-MS) analysis.

**Optical Spectroscopy**

**General** Fluorescent molecules were prepared as stock solutions in anhydrous DMSO and diluted such that the DMSO content did not exceed 1% (v/v) in assays. Fluorescence and absorbance measurements were performed on a plate reader (Spark® 20M, Tecan) equipped with filters and a monochromator, while extinction coefficient determination was carried out on a V-770 Spectrophotometer (Jasco) using quartz cuvettes. Bioluminescence spectra for *in vitro* measurements were obtained using the bioluminescence scan mode of a Cytation 5 plate reader (BioTeck), while cellular data was obtained on a Spark® 20M plate reader (Tecan). Absolute Quantum yields were determined using a Quantaurus-QY spectrometer (model C11374) from Hamamatsu with diluted samples (A ≈ 0.1). Results were processed and illustrated using Origin software. All reported values are averages of independent measurements or wells (n ≥ 3).

**Fluorescence/absorbance measurements.** Calcium dyes (100 µM) were pre-incubated with 400 µM protein (either HaloTag7 or BSA) for 2 h at room temperature (rt) in 1x PBS. Subsequently, the dye-protein mixture was diluted to a final dye concentration of 1 µM in 100 µL of a calcium buffer at varying concentrations (0-39 µM or 0-10’000 µM) in non-binding black flat-bottom 96-well plates (PerkinElmer). Calcium concentrations of buffers for $MaPCa_{high}$ experiments were adjusted by mixing different proportions of buffers from a commercially available kit (Invitrogen) following the vendors protocol [EGTA buffer (30 mM MOPS, 10 mM EGTA, 100 mM KCl, pH 7.2); Ca-EGTA buffer (30 mM MOPS, 10 mM Ca-EGTA, 100 mM KCl, pH 7.2]. Calcium concentrations of buffers for $MaPCa_{low}$ experiments were obtained by self-made buffer solutions (100 mM KCl, 30 mM

MOPS, pH 7.2) containing various calcium concentrations (0 µM, 1.35 µM, 2.85 µM, 10 µM, 20 µM, 50 µM, 100 µM, 350 µM, 1’000 µM, 3’500 µM, 10’000 µM). Fluorescence intensities were measured using a plate reader (Spark 20M, Tecan) after 30 min of incubation at rt with slow shaking (spectra were recorded with an excitation at 520 nm (TMR-variants), 570 nm (CPY-variants) or 610 nm (SiR-variants); band width (BW): 10 nm, step size: 1 nm). Absorbance measurements were performed analogously but in non-binding flat bottom transparent 96-well plates (ThermoScientific), at 5 µM final dye concentration (BW: 1 nm).

$K_D(Ca^{2+})$-values were obtained using linear regression of log [(F - Fmin)/(Fmax - F)] vs. log $[Ca^{2+}]$, where x-intercept= log $K_D(Ca^{2+})$.

**Bioluminescence measurements.** H-Luc protein (5 µM) was labeled with excess dye (20 µM) for at least 30 min at rt. The dye-protein mixture was diluted to 1 nM final protein concentration in 100 µL calcium buffer as previously described (see fluorescence measurements) supplemented with 5 mg/mL BSA. Before measurements, 2 µL of diluted NanoGlow furimazine solution (Promega) was spiked in the mixture (final concentrations: H-Luc: 1 nM, calcium indicator: 4 nM, NanoGlow: 800x, BSA: 5 mg/mL). Luminescence emission profile of each well were recorded on a microplate reader (Spark20M, Tecan) over the range 390-660 nm (18 measurements every 15 nm, BW 25 nm). For the photos of Figure 1, the final H-Luc concentration was at 10 nM with 40 nM dye, 400x NanoGlow and 5 mg/mL BSA.

**Cell culture and stable cell line generation**

**Flp-In 293 cells.** Flp-In 293 cells were maintained in high-glucose DMEM (Life Technologies) medium supplemented with GlutaMAX (Life Technologies), sodium pyruvate (Life Technologies), 10% FBS (Life Technologies) and phenol red in a humidified incubator at 37°C and 5% $CO_2$ atmosphere. Cells were passaged every 2-3 days or at confluency using 0.05% EGTA-free trypsin solution (Life Technologies). Cell lines were regularly tested for mycoplasma contamination *via* PCR. For live-cell imaging, cells were kept in fully supplemented high-glucose DMEM without additional phenol red and at 37°C and 5% $CO_2$ atmosphere.

Stable Flp-In T-Rex 293 cell lines were generated from commercially available cell lines (ThermoFisher Scientific, Catalog number: R78007) as previously described.[5] The Flp-In System was used to express a HaloTag-SNAP or H-Luc construct in different compartments (cytosol, nucleus) under control of the Cytomegalovirus (CMV) promoter. In brief, the protein encoding pcDNA5-FRT plasmids (without localization sequence (cyto) or with nuclear localization sequence (NLS, nucleus) and pOG44 that encodes Flippase recombinase were co-transfected into the Flp-In T-REx 293 cell line using Lipofectamine3000 transfection reagent (Life Technologies) following manufacturer's protocol. Homologous recombination of the FRT sites and the host cell chromosome yielded the stable cell lines. Selection was performed using 100 µg/mL Hygromycin B (ThermoFisher Scientific) and 15 µg/mL blasticidine (ThermoFisher Scientific).

##### Rat hippocampal neurons

**General** 24-well glass-bottom plates were coated with poly-L-ornithine (100 µg/mL) for 20 minutes, washed three times with 1x PBS and coated with laminin dissolved in 1x HBSS (1 µg/mL) for 1 hour. New born pups (0-1 days, WISTAR rats) were sacrificed and the hippocampi were isolated. Tryptic digest was followed by mechanical dissection using a pipette to obtain a homogenous solution. The solution was filtered through a cell strainer (40 µm pore size) and cell numbers were measured considering live and dead cells. Final cell numbers were adjusted to 55'000 live cells per well. 2 h after seeding, medium was removed and fresh phenol-red free neurobasal medium (NB) supplemented with antibiotics (Penicilin/Streptomycin, Life Technologies), GlutaMAX and B27 was added. Neurons were maintained in a humidified incubator at 37°C and 5% CO2 atmosphere. A third of the medium was exchanged every 5-8 days.

**rAAV production and neuron transduction.** Recombinant AAVs (rAAVs) were generated as described in Zolotukhin *et al.* (2002).[10] In brief, plasmids pRV1 (AAV2 Rep and Cap sequences), pH21 (AAV1 Rep and Cap sequences), pFD6 (Adenovirus helper plasmid) and the AAV plasmid containing the recombinant expression cassette flanked by AAV2 packaging signals (ITRs) were transfected via Polyethylenimine 25000 (PEI25000) into HEK293 cells. 5 days post transfection, the medium and cells were harvested. The cells were lysed using TNT extraction buffer (20 mM Tris pH7.5, 150 mM NaCl, 1% TX-100, 10 mM MgCl2). The cell debris was spun down and the cell supernatant treated with Benzonase. The rAAVs were purified from the medium and cell supernatant via FPLC using AVB Sepharose columns, which were subsequently concentrated using centrifugal filter devices (Amicon, Merck KGaA, Darmstadt, Germany) with a MWCO of 100 kDa. rAVV1 GCaMP6f are commercially available (Addgene 100837- AAV1).

The rAAV titers were evaluated by qPCR and are summarized in Supporting Table 3.

Hippocampal neurons were transduced with rAVVs after 7-8 days in culture. 0.5 µL of the respective rAAV were diluted into 10 µL of phenol-red free NB medium and subsequently added to corresponding samples. After the 13-18 days *in vitro* (DIV), the neurons were labelled with dyes as described for Flp-In 293 cells.

**Signal/background measurements.** Primary rat hippocampal neurons were transduced with a rAAV encoding a cytosolic (NES) HaloTag-eGFP while others were left non-transduced. All wells were labeled on DIV 14-18 with 1 µM $MaPCa_{high}$ AM-indicator and in presence of 0.04% Pluronic-F127 for at least 2 h. Transduced and untransduced neurons were subsequently imaged as previously explained for Flp-In 293 cells and data treatment was performed analogously.

**Protein Sequences**

Protein Amino Acid sequence of NES-HaloTag7-mEGFP expressed in cultured neurons

MLQNELALKLAGLDINKTGGSGSEIGTGFPFDPHYVEVLGERMHYVDVGPRDGTPVLFLHGNP
TSSYVWRNIIPHVAPTHRCIAPDLIGMGKSDKPDLGYFFDDHVRFMDAFIEALGLEEVVLVIHDW
GSALGFHWAKRNPERVKGIAFMEFIRPIPTWDEWPEFARETFQAFRTTDVGRKLIIDQNVFIEGT
LPMGVVRPLTEVEMDHYREPFLNPVDREPLWRFPNELPIAGEPANIVALVEEYMDWLHQSPVP
KLLFWGTPGVLIPPAEAARLAKSLPNCKAVDIGPGLNLLQEDNPDLIGSEIARWLSTLEISGMVSK
GEELFTGVVPILVELDGDVNGHKFSVSGEGEGDATYGKLTLKFICTTGKLPVPWPTLVTTLTYGV
QCFSRYPDHMKQHDFFKSAMPEGYVQERTIFFKDDGNYKTRAEVKFEGDTLVNRIELKGIDFKE
DGNILGHKLEYNYNSHNVYIMADKQKNGIKVNFKIRHNIEDGSVQLADHYQQNTPIGDGPVLLPD
NHYLSTQSKLSKDPNEKRDHMVLLEFVTAAGITLGMDELYK

Protein Amino Acid sequence of CalR-HaloTag7-P30-SNAP-KDEL expressed in cultured neurons

MLLSVPLLLGLLGLAVAGGSGGSEFGSEIGTGFPFDPHYVEVLGERMHYVDVGPRDGTPVLFL
HGNPTSSYVWRNIIPHVAPTHRCIAPDLIGMGKSDKPDLGYFFDDHVRFMDAFIEALGLEEVVLV
IHDWGSALGFHWAKRNPERVKGIAFMEFIRPIPTWDEWPEFARETFQAFRTTDVGRKLIIDQNV
FIEGTLPMGVVRPLTEVEMDHYREPFLNPVDREPLWRFPNELPIAGEPANIVALVEEYMDWLHQ
SPVPKLLFWGTPGVLIPPAEAARLAKSLPNCKAVDIGPGLNLLQEDNPDLIGSEIARWLSTLEISG
SGRPPPPPPPPPPPPPPPPPPPPPPPPPPPPPPGGRSRSLEMDKDCEMKRTTLDSPLGKLELS
GCEQGLHEIIFLGKGTSAADAVEVPAPAAVLGGPEPLMQATAWLNAYFHQPEAIEEFPVPALHH
PVFQQESFTRQVLWKLLKVVKFGEVISYSHLAALAGNPAATAAVKTALSGNPVPILIPCHRVVQG
DLDVGGYEGGLAVKEWLLAHEGHRLGKPGLGGGGGSGKDEL

Protein Amino Acid sequence of HaloTag7-P30-SNAP-3xNLS expressed in mammalian cells

MGSEIGTGFPFDPHYVEVLGERMHYVDVGPRDGTPVLFLHGNPTSSYVWRNIIPHVAPTHRCIA
PDLIGMGKSDKPDLGYFFDDHVRFMDAFIEALGLEEVVLVIHDWGSALGFHWAKRNPERVKGIA
FMEFIRPIPTWDEWPEFARETFQAFRTTDVGRKLIIDQNVFIEGTLPMGVVRPLTEVEMDHYREP
FLNPVDREPLWRFPNELPIAGEPANIVALVEEYMDWLHQSPVPKLLFWGTPGVLIPPAEAARLA
KSLPNCKAVDIGPGLNLLQEDNPDLIGSEIARWLSTLEISGSGRPPPPPPPPPPPPPPPPPPPP
PPPPPPPPPGGRSRSLEMDKDCEMKRTTLDSPLGKLELSGCEQGLHEIIFLGKGTSAADAVEVP
APAAVLGGPEPLMQATAWLNAYFHQPEAIEEFPVPALHHPVFQQESFTRQVLWKLLKVVKFGE
VISYSHLAALAGNPAATAAVKTALSGNPVPILIPCHRVVQGDLDVGGYEGGLAVKEWLLAHEGH
RLGKPGLGGAPDPKKKRKVDPKKKRKVDPKKKRKELRASPQ

Protein Amino Acid sequence of HLuc-NLS expressed in mammalian cells

MGSEIGTGFPFDPHYVEVLGERMHYVDVGPRDGTPVLFLHGNPTSSYVWRNIIPHVAPTHRCIA
PDLIGMGKSDKPDLGYFFDDHVRFMDAFIEALGLEEVVLVIHDWGSALGFHWAKRNPERVKGIA
FMEFIRPIPTWDEWPEFARETFQAFRTSGDQMGQIEKIFKVVYPVDDHHFKVILHYGTLVIDGVT
PNMIDYFGRPYEGIAVFDGKKITVTGTLWNGNKIIDERLINPDGSLLFRVTINGVTGWRLCERILA
GGTGGSGGTGGSMVFTLEDFVGDWRQTAGYNLDQVLEQGGVSSLFQNLGVSVTPIQRIVLSG

ENGLKIDIHVIIPYEGLDVGRKLIIDQNVFIEGTLPMGVVRPLTEVEMDHYREPFLNPVDREPLWR
FPNELPIAGEPANIVALVEEYMDWLHQSPVPKLLFWGTPGVLIPPAEAARLAKSLPNCKAVDIGP
GLNLLQEDNPDLIGSEIARWLSTLEISGKSGLRSRADPKKKRKVDPKKKRKVDPKKKRKVGSTG
SR

Color code: Localizations - HaloTag7 - HLuc - SNAP-Tag - T2A - linker - mEGFP

### Chemical synthesis and characterization

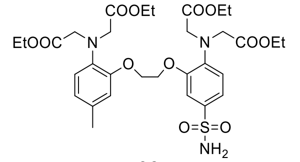

**03**

A mixture of thionyl chloride (169 µL, 2.32 mmol, 7.0 eq.) and chlorosulfonic acid (122 µL, 1.83 mmol, 5.5 eq.) was cooled on an ice bath for 15 minutes. Then, a solution of N-[2-[2-[2-[Bis(2-ethoxy-2-oxoethyl)amino]-5-methylphenoxy]ethoxy]phenyl]-N-(2-ethoxy-2-oxoethyl)glycine ethyl ester* **01** (200mg, 332 µmol, 1.0 eq.) in DCM (1.6 mL) was added slowly. The solution was allowed to warm to room temperature and stirred for 24 h. The solution was slowly added to a cooled mixture of EtOAc and aqueous ammonia (≈ 10 mL). The resulting mixture was further stirred for 2 h. The product was extracted with EtOAc and the combined organic layers dried over magnesium sulfate, filtered and concentrated under reduced pressure. The crude product was purified by flash column chromatography (25 g SiO2 column, 20-60% EtOAc in hexanes) to give **03** as an off-white powder (141 mg, 62%). $^{1}H$ NMR (400 MHz, $CDCl_3$): δ = 7.46 (s, 1H), 6.93 – 6.79 (m, 4H), 6.74 (s, 1H), 4.92 (d, J = 4.5 Hz, 2H), 4.30 (hept, J = 2.7 Hz, 4H), 4.11 (d, J = 6.4 Hz, 8H), 4.03 (p, J = 7.0 Hz, 8H), 2.57 (s, 3H), 1.15 (td, J = 7.1, 4.2 Hz, 12H); $^{13}C$ NMR (100 MHz, $CDCl_3$): δ = 171.67, 171.29, 153.16, 150.25, 139.53, 137.02, 132.15, 131.20, 122.41, 121.89, 119.32, 119.28, 116.14, 113.62, 67.52, 66.98, 61.15, 60.93, 53.59, 53.49, 20.00, 14.17, 14.13; HRMS (ESI$^+$) m/z calcd. for $C_{31}H_{43}N_3O_{12}S$ [M]+, 682.2640; found 682.2635.

* synthesized according to published procedures[11]

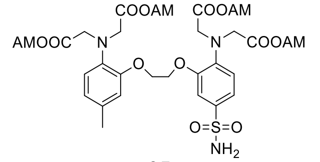

**05**

Sulfonamide **03** (110 mg, 161 µmol, 1 eq.) was dissolved in anhydrous DCM (1.1 mL). Then 4-dimethylaminopyridine (29.6 mg, 242 µmol, 1.5 eq.) and di-tert-butyl dicarbonate (276 µL, 1.29 mmol, 8.0 eq.) were added and the solution was stirred for 24 h at 35°C. The product was extracted with DCM and the combined organic layers dried over magnesium sulfate, filtered and concentrated under reduced pressure. The crude was purified by flash column chromatography

(25 g SiO2 column, 20-70% EtOAc in hexanes) to give Boc-protected **03** as an off-white powder (80 mg, 59%). It was directly re-dissolved in methanol (500 µL) and tetrahydrofuran (4 mL). Subsequently, potassium hydroxide (1 M, 382 µL, 382 µmol, 4 eq.) was added followed by potassium hydroxide (0.1M, 2.86 mL, 286 µmol, 3 eq.). The solution was stirred at room temperature for 2 h. In the following, the solution was neutralized using 0.1 M HCl and acetic acid. The mixture was dried by lyophilization and the crude re-suspended in MeCN (2 mL). Then add DIPEA (316 µL, 1.91 mmol, 20 eq.) and bromomethyl acetate (65.5 µL, 668 µmol, 7 eq.) was added. After 48 h, the solvents were evaporated and the crude residue was suspended in DCM (2 mL) and TFA (200 µL). The resulting solution was stirred for 2 h before being neutralized using $Na_2CO_3$. The product was extracted with EtOAc and the combined organic layers were dried over magnesium sulfate, filtered and concentrated under reduced pressure. The crude mixture was purified by HPLC (50 mL/min; A/B 30-80%; 60 min) to obtain **05** as white powder (48 mg, 61%). $^1$H NMR (400 MHz, $CDCl_3$) δ = 7.43 (d, J = 2.4 Hz, 1H), 7.01 – 6.84 (m, 4H), 6.77 (s, 1H), 5.58 (d, J = 2.8 Hz, 12H), 5.19 (s, 2H), 4.33 (dq, J = 7.6, 4.3, 3.5 Hz, 4H), 4.16 (d, J = 4.4 Hz, 8H), 2.60 (d, J = 2.5 Hz, 3H), 2.16 – 1.99 (m, 12H); $^{13}$C NMR (101 MHz, $CDCl_3$) δ = 170.32, 170.28, 169.85, 169.82, 153.20, 150.51, 138.79, 135.98, 132.72, 132.07, 123.26, 122.04, 120.10, 120.07, 116.20, 113.61, 79.46, 79.34, 77.48, 77.36, 77.16, 76.84, 67.43, 66.97, 53.55, 53.38, 20.86, 20.84, 20.17. HRMS (ESI$^+$) m/z calcd. for $C_{35}H_{43}N_3O_{20}S$ [M+Na]$^+$, 880.2053; found 880.2052.

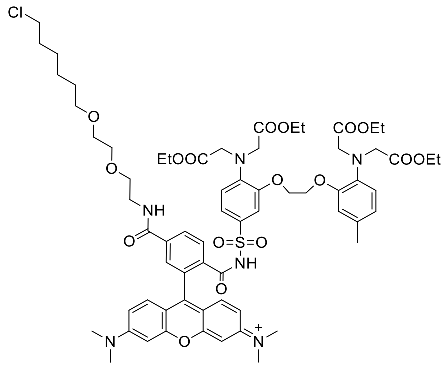

**S02**

TMR-CA (1.00 mg, 1.57 µmol, 1.0 eq.) and sulfonamide **03** (1.29 mg, 1.89 µmol, 1.2 eq.) were solubilized in DCM (1 mL). Then, DMAP (1.54 mg, 12.6 µmol, 8.0 eq.) and EDC-HCl (2.41 mg, 12.6 µmol, 8.0 eq.) were added and the tube was sealed. The solution was stirred at 60°C overnight. The crude mixture was purified by HPLC (4 mL/min; A/B 30-90%; 60 min) to obtain rhodamine conjugate **S02** as red powder (1.20 mg, 56%). $^1$H NMR (400 MHz, $CDCl_3$): δ = 7.96 –

7.91 (m, 2H), 7.55 – 7.50 (m, 1H), 7.39 (s, 1H), 7.03 – 6.74 (m, 5H), 6.67 – 6.57 (m, 5H), 4.30 (s, 4H), 4.11 – 3.93 (m, 16H), 3.70 – 3.56 (m, 4H), 3.55 – 3.45 (m, 3H), 3.40 (t, *J* = 6.7 Hz, 2H), 2.31 (s, 3H), 1.81 – 1.66 (m, 2H), 1.54 (p, *J* = 6.8 Hz, 2H), 1.47 – 1.37 (m, 2H), 1.36 – 1.27 (m, 2H), 1.13 (td, *J* = 7.1, 3.8 Hz, 12H). $^{13}$C NMR (100 MHz, $CDCl_3$): δ = 171.85, 171.44, 166.45, 165.58, 154.05, 153.89, 150.41, 139.61, 139.05, 136.24, 129.68, 135.01, 129.50, 125.93, 122.03, 119.42, 113.61, 111.32, 98.64, 77.48, 77.16, 76.85, 71.35, 70.35, 70.09, 69.66, 67.40, 61.36, 61.24, 54.18, 45.20, 40.96, 40.18, 32.65, 29.45, 26.80, 25.50, 20.16, 14.16, 14.12. HRMS (ESI$^+$) m/z calcd. for $C_{66}H_{83}ClN_6O_{17}S$ $[M]^{2+}$, 650.2685; found 650.2680.

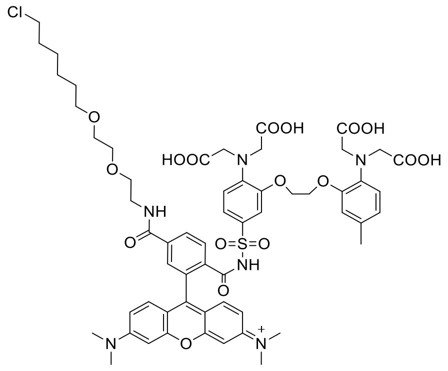

**MaPCa-558$_{high}$**

To a solution of **S02** (500 µg, 0.348 µmol, 1.0 eq.) in MeOH (100 µL) and THF (100 µL) was added an aqueous solution of potassium hydroxide (0.1M, 192 µL, 19.2 µmol, 50 eq.) and the resulting solution stirred for 6 h at rt. In the following, the solution was neutralized by the addition of 0.1 M HCl and acetic acid. The solvents were evaporated and the crude mixture purified by HPLC (4 mL/min; A/B 10-90%; 60 min) to obtain **MaPCa-558$_{high}$** as red powder (0.30 mg, 66%). $^1$H NMR (400 MHz, DMSO): δ= 12.39 (s, 4H), 8.69 (t, *J* = 5.7 Hz, 1H), 8.02 (d, *J* = 8.1 Hz, 1H), 7.91 (d, *J* = 7.9 Hz, 1H), 7.42 (s, 1H), 7.01 – 6.92 (m, 2H), 6.90 – 6.70 (m, 4H), 6.50 – 6.36 (m, 6H), 4.27 (dq, *J* = 9.9, 5.6 Hz, 4H), 3.97 (d, *J* = 32.2 Hz, 8H), 3.58 (t, *J* = 6.6 Hz, 2H), 3.44 (s, 6H), 3.28 (d, *J* = 6.5 Hz, 4H), 2.94 (s, 12H), 2.21 (s, 3H), 1.66 (p, *J* = 6.7 Hz, 2H), 1.40 (p, *J* = 7.1 Hz, 2H), 1.37 – 1.24 (m, 2H), 1.24 (dq, *J* = 8.9, 4.5, 3.9 Hz, 2H).$^{13}$C NMR (101 MHz, DMSO) δ 172.49, 171.86, 165.41, 164.66, 153.35, 152.34, 151.29, 149.43, 140.27, 139.45, 136.63, 133.27, 128.77, 128.64, 121.72, 121.12, 118.23, 115.19, 108.49, 106.27, 98.20, 70.09, 69.51, 69.34, 69.05, 68.63, 67.16, 53.51, 52.89, 45.35, 40.14, 39.94, 39.73, 39.52, 39.31, 39.10, 38.89, 31.99, 28.97, 26.07, 24.87, 19.61. HRMS (ESI$^+$) m/z calcd. for $C_{58}H_{68}N_6O_{17}ClS$ $[M]^{2+}$, 594.2059; found 594.2057.

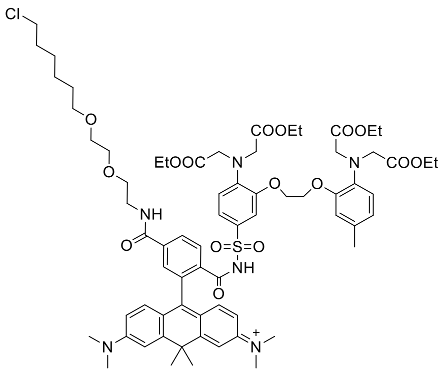

**S03**

To a solution of CPY-CA (5.00 mg, 7.54 µmol, 1.0 eq.) in anhydrous DCM (1.5 mL) was added thionyl chloride (8.20 µL, 113 µmol, 15 eq.) followed by pyridine (4.88 µL, 60.3 µmol, 8.0 eq.). The solution was stirred for 30 minutes at room temperature. In the following, the mixture was heated to 60°C until most of the solvent had evaporated. Then, the vial was closed and the crude dried under high vacuum. After 1 h, a solution of BAPTA-Et-sulfonamide **03** (6.17 mg, 9.05 µmol, 1.2 eq.), DIPEA (37.4 µL, 226 µmol, 30 eq.) and 4-dimethylaminopyridine (1.84 mg, 15.1 µmol, 2.0 eq.) in anhydrous MeCN (2.38 mL) was added under argon atmosphere. The solution was heated to 60°C and stirred for 1 h. The solvents were evaporated and the crude mixture purified by HPLC (8 mL/min; A/B 50-90%; 60 min) to obtain **S04** as blue powder (4.20 mg, 42%). $^{1}$H NMR (400 MHz, $CDCl_3$): δ = 7.92 (d, J = 8.0 Hz, 1H), 7.75 (dd, J = 8.0, 1.4 Hz, 1H), 7.48 (d, J = 2.5 Hz, 2H), 7.39 (s, 1H), 7.08 (d, J = 1.4 Hz, 1H), 6.96 – 6.83 (m, 6H), 6.74 (d, J = 8.8 Hz, 2H), 6.58 (s, 1H), 4.31 (s, 4H), 4.09 – 3.95 (m, 16H), 3.61 – 3.47 (m, 10H), 3.39 (t, J = 6.7 Hz, 2H), 3.12 (s, 12H), 2.11 (s, 3H), 1.91 (d, J = 3.9 Hz, 6H), 1.73 (dt, J = 14.7, 6.8 Hz, 2H), 1.53 (p, J = 6.9 Hz, 2H), 1.47 – 1.36 (m, 2H), 1.36 – 1.27 (m, 2H), 1.13 (q, J = 7.0 Hz, 13H). $^{13}$C NMR (100 MHz, $CDCl_3$): δ = 171.78, 171.22, 166.59, 166.16, 161.86, 161.49, 154.26, 150.28, 146.48, 145.87, 141.09, 129.47, 129.14, 127.37, 124.76, 123.26, 122.00, 121.79, 119.30, 117.68, 115.97, 115.38, 77.48, 77.36, 77.16, 76.84, 71.32, 70.95, 70.28, 70.00, 69.59, 67.35, 61.33, 61.13, 53.91, 45.20, 43.78, 40.03, 38.42, 36.09, 33.25, 32.58, 29.43, 26.74, 25.45, 20.41, 14.16, 14.13, 1.17. HRMS ($ESI^+$) m/z calcd. for $C_{69}H_{89}ClN_6O_{16}S$ $[M]^{2+}$, 663.2945; found 663.2939.

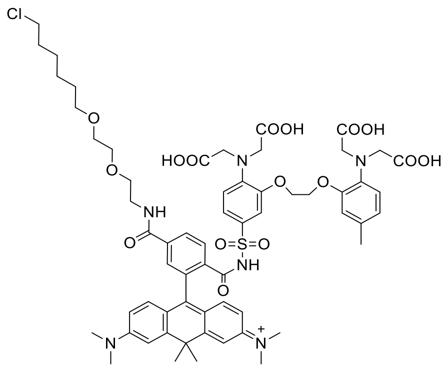

**MaPCa-619$_{high}$**

To a solution of **S03** (5.20 mg, 3.92 µmol, 1.0 eq.) in methanol (1.0 mL) and tetrahydrofuran (1.0 mL) was added potassium hydroxide (0.1M, 1.2 mL). The mixture was stirred at room temperature for 8 h. The solution was neutralized by the addition of 0.1 M HCl and acetic acid. The solvents were evaporated and the crude mixture purified by HPLC (4 mL/min; A/B 40-80%; 60 min) to obtain **MaPCa-619$_{high}$** as blue powder (2.00 mg, 42%). $^{1}H$ NMR (400 MHz, DMSO): δ= 8.67 (t, J = 5.8 Hz, 1H), 7.90 (d, J = 2.1 Hz, 2H), 7.14 (s, 1H), 7.04 – 6.79 (m, 5H), 6.79 – 6.71 (m, 2H), 6.53 (dd, J = 9.0, 2.5 Hz, 2H), 6.45 (dd, J = 8.9, 5.4 Hz, 2H), 4.33 – 4.24 (m, 4H), 3.99 (d, J = 27.8 Hz, 8H), 3.58 (t, J = 6.6 Hz, 2H), 3.40 (ddd, J = 17.1, 6.1, 3.5 Hz, 6H), 3.28 (t, J = 6.6 Hz, 2H), 2.94 (s, 12H), 2.11 (s, 3H), 1.85 (s, 6H), 1.65 (p, J = 6.7 Hz, 2H), 1.46 – 1.17 (m, 6H). $^{13}C$ NMR (100 MHz, DMSO): δ = 172.49, 171.91, 166.37, 164.89, 158.24, 157.90, 154.77, 153.17, 149.42, 144.93, 140.37, 139.46, 136.54, 133.14, 129.06, 128.31, 127.99, 127.47, 123.94, 122.34, 121.64, 121.07, 120.17, 118.20, 115.84, 115.28, 112.26, 71.93, 70.09, 69.51, 69.35, 68.57, 67.12, 53.50, 52.86, 45.34, 40.15, 39.94, 39.73, 39.52, 39.31, 39.10, 38.89, 37.59, 35.66, 32.78, 31.97, 28.98, 26.06, 24.86, 19.64. HRMS (ESI$^{+}$) m/z calcd. for $C_{61}H_{74}ClN_6O_{16}S$ $[M]^{2+}$, 607.2319; found 607.2311.

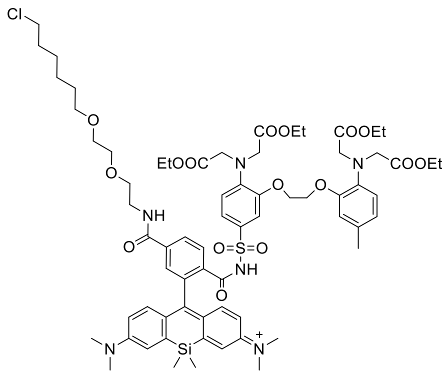

**S04**

To a solution of SiR-CA (5.00 mg, 7.36 µmol, 1.0 eq.) in anhydrous DCM (1.4 mL) was added thionyl chloride (8.10 µL, 110 µmol, 15 eq.) followed by pyridine (4.76 µL, 58.9 µmol, 8.0 eq.). The solution was stirred for 30 minutes at room temperature. In the following, the mixture was heated to 60°C until almost no solvent was left. Then, the vial was closed and the crude dried under high vacuum. After 1 h, a solution of BAPTA-Et-sulfonamide **03** (6.17 mg, 9.05 µmol, 1.2 eq.), DIPEA (37.4 µL, 226 µmol, 30 eq.) and 4-dimethylaminopyridine (1.84 mg, 15.1 µmol, 2.0 eq.) in anhydrous MeCN (2.4 mL) was added under argon atmosphere. The solution was heated to 60°C and stirred for 1 h. The solvents were evaporated and the crude mixture purified by HPLC (8 mL/min; A/B 50-90%; 60 min) to obtain **S04** as pale green-blue powder (4.20 mg, 41%). $^{1}$H NMR (400 MHz, $CDCl_3$): δ = 7.92 (d, J = 8.1 Hz, 1H), 7.72 (d, J = 9.4 Hz, 1H), 7.58 – 7.43 (m, 2H), 7.12 – 7.02 (m, 3H), 6.95 – 6.77 (m, 6H), 6.73 – 6.63 (m, 1H), 6.53 (s, 1H), 4.30 (s, 4H), 4.16 – 3.96 (m, 16H), 3.62 – 3.47 (m, 10H), 3.40 (t, J = 6.7 Hz, 2H), 3.10 (s, 12H), 1.97 (s, 3H), 1.80 – 1.68 (m, 2H), 1.54 (p, J = 6.8 Hz, 2H), 1.47 – 1.37 (m, 2H), 1.38 – 1.27 (m, 2H), 1.17 – 1.12 (m, 12H), 0.65 (d, J = 18.6 Hz, 6H). $^{13}$C NMR (100 MHz, $CDCl_3$): δ = 171.94, 166.98, 166.24, 156.11, 154.11, 150.44, 147.23, 140.96, 135.89, 134.10, 130.37, 129.71, 129.46, 126.89, 124.66, 122.89, 122.06, 121.96, 119.36, 117.94, 116.61, 115.31, 77.48, 77.36, 77.16, 76.84, 75.58, 71.33, 70.35, 70.08, 69.66, 67.26, 66.66, 61.49, 61.31, 54.31, 53.57, 45.17, 41.75, 40.02, 32.62, 29.85, 29.49, 26.77, 25.50, 22.84, 20.18, 14.16, 1.17, 0.12, -0.60. HRMS (ESI$^{+}$) m/z calcd. for $C_{68}H_{89}ClN_6O_{16}SSi$ $[M]^{2+}$, 671.2830; found 671.2821.

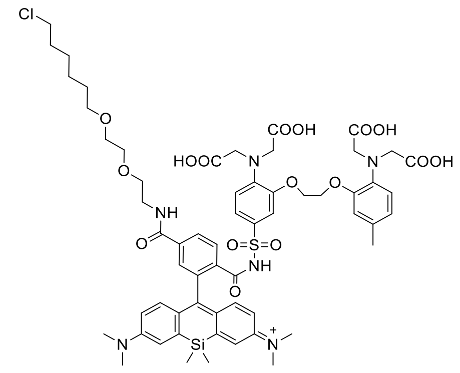

**MaPCa-656$_{high}$**

To a solution of **S04** (6.00 mg, 4.47 μmol, 1.0 eq.) in methanol (1.0 mL) and tetrahydrofuran (1.0 mL), was added potassium hydroxide (0.1M, 1.3 mL). The mixture was stirred at rt for 8 h. The solution was neutralized by the addition of 0.1 M HCl and acetic acid. The solvents were evaporated and the crude mixture purified by HPLC (4 mL/min; A/B 30-70%; 60 min) to obtain **MaPCa-656$_{high}$** as pale green-blue powder (3.10 mg, 56%). $^{1}$H NMR (400 MHz, DMSO): δ = 8.65 (t, J = 5.7 Hz, 1H), 7.95 – 7.84 (m, 2H), 7.22 (s, 1H), 7.04 – 6.92 (m, 4H), 6.85 (q, J = 6.6, 6.0 Hz, 2H), 6.79 – 6.69 (m, 2H), 6.59 (d, J = 9.1 Hz, 2H), 6.47 (d, J = 8.9 Hz, 2H), 4.26 (h, J = 5.9 Hz, 4H), 3.98 (d, J = 28.3 Hz, 8H), 3.58 (t, J = 6.6 Hz, 2H), 3.47 – 3.36 (m, 6H), 3.32 – 3.24 (m, 4H), 2.92 (s, 12H), 1.96 (s, 3H), 1.65 (p, J = 6.9 Hz, 2H), 1.46 – 1.37 (m, 2H), 1.37 – 1.24 (m, 2H), 1.25 – 1.15 (m, 2H), 0.56 (d, J = 21.8 Hz, 6H). $^{13}$C NMR (100 MHz, DMSO): δ =172.48, 171.93, 166.49, 164.86, 158.34, 157.97, 155.97, 153.13, 149.40, 148.05, 140.33, 139.44, 136.47, 134.52, 132.97, 129.31, 128.91, 128.28, 126.87, 125.32, 124.17, 122.27, 121.69, 121.07, 120.16, 118.20, 117.16, 115.09, 70.09, 69.51, 69.35, 68.58, 67.09, 53.49, 52.92, 45.35, 40.15, 39.94, 39.73, 39.52, 39.31, 39.10, 38.89, 31.97, 30.70, 28.98, 26.06, 24.86, 19.35, -0.08, -0.74. HRMS (ESI$^{+}$) m/z calcd. for $C_{60}H_{73}ClN_6O_{16}SSi$ $[M]^{2+}$, 615.2204; found 615.2196.

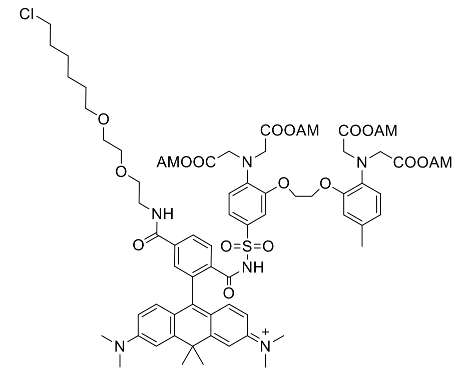

**MaPCa-619$_{high}$ AM**

To a solution of CPY-CA (5.00 mg, 7.54 µmol, 1.0 eq.) in anhydrous DCM (1.4 mL) was added thionyl chloride (8.20 µL, 113 µmol, 15 eq.) followed by pyridine (4.88 µL, 60.3 µmol, 8.0 eq.). The solution was stirred for 30 minutes. The mixture was heated to 60°C until almost no solvent was left. Then, the vial was closed and the crude dried under high vacuum. After 1 h, a solution of freshly prepared BAPTA-AM-sulfonamide **05** (7.76 mg, 9.05 µmol, 1.2 eq.), DIPEA (37.4 µL, 226 µmol, 30 eq.) and 4-dimethylaminopyridine (1.84 mg, 15.1 µmol, 2.0 eq.) in anhydrous MeCN (2.4 mL) was added under argon atmosphere. The solution was heated to 60°C and stirred for 1 h. The solvents were evaporated and the crude mixture purified by HPLC (8 mL/min; A/B 30-90%; 60 min) to obtain **MaPCa-619$_{high}$ AM** as blue powder (3.0 mg, 26%). $^{1}$H NMR (400 MHz, DMSO): δ = 8.68 (t, J = 5.6 Hz, 1H), 7.90 (s, 2H), 7.13 (s, 1H), 7.02 (s, 1H), 6.97 – 6.94 (m, 2H), 6.87 (dtd, J = 16.8, 7.4, 1.7 Hz, 2H), 6.80 (s, 1H), 6.73 (dd, J = 7.7, 1.9 Hz, 1H), 6.53 (dd, J = 9.0, 2.5 Hz, 2H), 6.41 (d, J = 8.8 Hz, 2H), 5.54 (d, J = 2.3 Hz, 8H), 4.25 (dd, J = 15.6, 5.0 Hz, 4H), 4.11 (d, J = 15.4 Hz, 8H), 3.57 (t, J = 6.6 Hz, 2H), 3.47 – 3.35 (m, 8H), 3.28 (t, J = 6.5 Hz, 2H), 2.95 (s, 12H), 2.04 (s, 3H), 1.98 (d, J = 5.3 Hz, 12H), 1.85 (d, J = 5.7 Hz, 6H), 1.70 – 1.61 (m, 2H), 1.40 (p, J = 6.7 Hz, 2H), 1.36 – 1.27 (m, 2H), 1.31 – 1.17 (m, 2H). $^{13}$C NMR (100 MHz, $CDCl_3$): δ = 169.77, 169.20, 166.33, 164.86, 158.31, 157.96, 155.45, 153.32, 149.63, 149.49, 144.98, 140.44, 138.33, 135.40, 133.89, 129.00, 128.27, 127.88, 127.26, 123.60, 122.25, 121.84, 121.32, 120.40, 118.49, 117.60, 115.61, 114.64, 113.95, 112.31, 110.54, 110.25, 78.97, 71.91, 70.09, 69.52, 69.36, 68.57, 53.05, 52.63, 45.35, 40.31, 40.14, 39.94, 39.73, 39.52, 39.31, 39.10, 38.89, 37.60, 35.63, 32.69, 31.97, 28.98, 26.06, 24.86, 20.40, 19.45. HRMS (ESI$^+$) m/z calcd. for $C_{73}H_{89}ClN_6O_{24}S$ $[M]^{2+}$, 751.2741; found 751.2738.

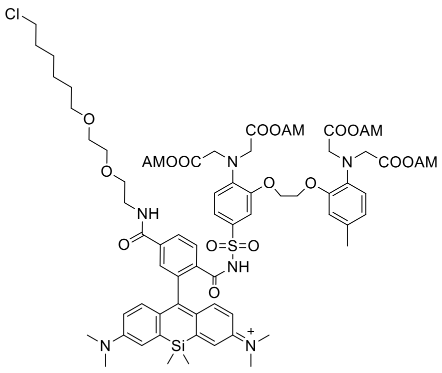

**MaPCa-656$_{high}$ AM**

To a solution of SiR-CA (5.00 mg, 7.36 µmol, 1.0 eq.) in anhydrous DCM (1.4 mL) was added thionyl chloride (8.00 µL, 110 µmol, 15 eq.) followed by pyridine (4.76 µL, 58.9 µmol, 8.0 eq.). The solution was stirred for 30 minutes. In the following, the mixture was heated to 60°C until almost no solvent was left. Then, the vial was closed and the crude dried under high vacuum. After 1 h, a solution of freshly prepared BAPTA-AM-sulfonamide **05** (7.58 mg, 8.83 µmol, 1.2 eq.), DIPEA (36.5 µL 221 µmol, 30 eq.) and 4-dimethylaminopyridine (1.80 mg, 14.7 µmol, 2.0 eq.) in anhydrous MeCN (3.1 mL) was added under argon atmosphere. The solution was heated to 60°C and stirred for 1 h. The solvents were evaporated and the crude mixture purified by HPLC (8 mL/min; A/B 30-90%; 60 mins) to obtain **MaPCa-656$_{high}$ AM** as pale green-blue powder (4.30 mg, 38%). $^1$H NMR (400 MHz, $CDCl_3$): δ = 7.94 (d, J = 8.0 Hz, 1H), 7.73 (dd, J = 8.0, 1.4 Hz, 1H), 7.46 – 7.41 (m, 2H), 7.11 (d, J = 1.2 Hz, 1H), 7.04 (dd, J = 9.0, 2.8 Hz, 2H), 7.00 – 6.86 (m, 4H), 6.82 (d, J = 9.0 Hz, 2H), 6.65 (t, J = 5.3 Hz, 1H), 6.58 (s, 1H), 5.61 (d, J = 12.3 Hz, 8H), 4.33 (s, 4H), 4.14 (d, J = 12.9 Hz, 8H), 3.62 – 3.47 (m, 10H), 3.41 (t, J = 6.7 Hz, 2H), 3.12 (s, 12H), 2.04 (d, J = 10.0 Hz, 12H), 1.99 (s, 3H), 1.74 (dt, *J* = 14.6, 6.7 Hz, 2H), 1.54 (p, J = 6.8 Hz, 2H), 1.48 – 1.38 (m, 2H), 1.37 – 1.27 (m, 2H), 0.64 (d, J = 15.8 Hz, 6H). $^{13}$C NMR (100 MHz, $CDCl_3$): δ = 170.34, 170.33, 169.87, 169.80, 169.67, 169.66, 166.71, 166.11, 154.85, 154.43, 150.58, 144.99, 141.03, 138.75, 136.36, 135.89, 130.83, 129.13, 127.05, 125.12, 123.57, 123.48, 122.24, 121.43, 120.34, 119.43, 115.64, 114.05, 79.50, 79.46, 77.48, 77.36, 77.16, 76.84, 74.72, 71.36, 70.34, 70.04, 69.56, 67.40, 66.96, 53.67, 53.48, 45.17, 43.90, 40.12, 32.60, 29.46, 26.75, 25.47, 20.80, 20.77, 20.17, 1.17, -0.07, -0.77. HRMS (ESI$^+$) m/z calcd. for $C_{72}H_{89}ClN_6O_{24}SSi$ $[M]^{2+}$, 759.2626; found 759.2618.

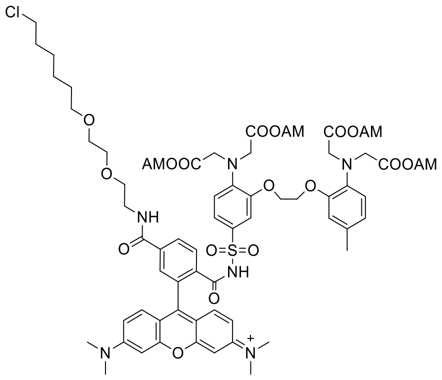

**MaPCa-558$_{high}$ AM**

To a solution of TMR-CA (3.50 mg, 5.50 µmol, 1.0 eq.) in anhydrous DCM (1.0 mL) was added thionyl chloride (6.00 µL, 83.0 µmol, 15 eq.) followed by pyridine (3.56 µL, 44.0 µmol, 8.0 eq.). The solution was stirred for 30 minutes. Upon complete activation, the mixture was heated to 60°C until almost no solvent was left. Then, the vial was closed and the crude dried under high vacuum. After 1 h, a solution of freshly prepared BAPTA-AM-sulfonamide **05** (5.00 mg, 5.80 µmol, 1.1 eq.), DIPEA (27.0 µL 165 µmol, 30 eq.) and 4-dimethylaminopyridine (1.34 mg, 11.0 µmol, 2.0 eq.) in anhydrous MeCN (2.3 mL) was added under argon atmosphere. The solution was heated to 60°C and stirred for 1 h. The solvents were evaporated and the crude mixture purified by HPLC (8 mL/min; A/B 30-80%; 60 min) to obtain **MaPCa-558$_{high}$ AM** as red powder (2.30 mg, 28%). $^{1}$H NMR (400 MHz, DMSO): δ = 8.70 (t, J = 5.5 Hz, 1H), 8.04 (d, J = 8.1 Hz, 1H), 7.91 (d, J = 8.0 Hz, 1H), 7.44 (s, 1H), 7.02 – 6.93 (m, 2H), 6.93 – 6.80 (m, 3H), 6.74 (dd, J = 7.7, 1.9 Hz, 1H), 6.43 (d, J = 16.6 Hz, 6H), 5.53 (s, 8H), 4.26 (h, J = 3.5 Hz, 4H), 4.10 (d, J = 20.7 Hz, 8H), 3.58 (t, J = 6.6 Hz, 2H), 3.49 – 3.38 (m, 4H), 3.29 (t, J = 6.5 Hz, 2H), 2.95 (s, 12H), 2.17 (s, 3H), 1.98 (s, 12H), 1.71 – 1.60 (m, 2H), 1.46 – 1.36 (m, 2H), 1.36 – 1.28 (m, 2H), 1.30 – 1.17 (m, 2H). $^{13}$C NMR (100 MHz, DMSO): δ = 169.75, 169.24, 169.22, 169.12, 165.45, 164.63, 158.25, 157.91, 153.60, 152.37, 151.33, 149.67, 140.22, 138.32, 135.36, 128.61, 128.31, 121.89, 121.35, 118.56, 115.68, 114.04, 108.53, 106.49, 98.25, 78.97, 78.86, 70.10, 69.53, 69.36, 68.64, 67.17, 66.83, 53.04, 52.63, 45.36, 40.15, 39.94, 39.73, 39.52, 39.31, 39.10, 38.89, 32.00, 28.99, 26.08, 24.88, 24.85, 20.43, 20.40, 19.49. HRMS (ESI$^{+}$) m/z calcd. for $C_{70}H_{83}ClN_6O_{25}S$ $[M]^{2+}$, 738.2481; found 738.2479.

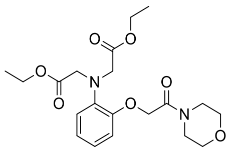

**02**

2-(2-morpholin-4-yl-2-oxoethoxy)aniline **S01** (300 mg, 1.27 mmol, 1.0 eq.) was dissolved in acetonitrile (12 mL). Then sodium iodide (952 mg, 6.35 mmol, 5.0 eq.), DIPEA (1.05 mL, 6.35 mmol, 5.0 eq.) and ethyl bromoacetate (704 µL, 6.35 mmol, 5.0 eq.) were added. The reaction mixture was stirred at 100°C for 20 h and then quenched by addition of water. The mixture was extracted with DCM (3x), combined organic phases were dried over magnesium sulfate and the solvent was removed *in vacuo*. The crude was purified by flash column chromatography (25 g $SiO_2$, 20 - 60% EtOAc in hexane) and the pure product **02** was obtained as a beige solid (511 mg, 98%).$^{1}$H NMR (400 MHz, $CDCl_3$): δ = 6.93 (s, 4H), 4.74 (s, 2H), 4.15 (q, $J$ = 7.1 Hz, 4H), 4.13 (s, 4H), 3.64 – 3.62 (m, 4H), 3.58 – 3.59 (m, 4H), 1.25 (t, $J$ = 7.1 Hz, 6H); $^{13}$C NMR (101 MHz, $CDCl_3$): δ 171.3, 166.9, 150.2, 140.2, 139.7, 123.0, 122.8, 120.4, 115.5, 68.6, 67.0, 60.8, 53.8, 14.4; HRMS (ESI$^+$) m/z calcd. for $C_{20}H_{28}N_2O_7$ $[M+H]^+$ 409.1969, found 409.1967.

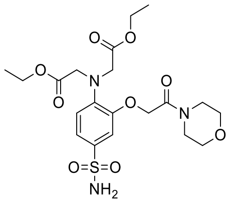

**04**

MOBHA-Et **02** (511 mg, 1.25 mmol, 1.0 eq.) was dissolved in dichloromethane (5.8 mL) and added to pre-cooled (0°C) chlorosulfonic acid (5.83 mL, 87.5 mmol, 70 eq.). The reaction mixture was stirred at 0°C for 30 min. The cooling bath was removed and the reaction mixture was stirred at rt for further 18.5 h. The mixture was diluted with EtOAc (20 mL) and cooled to 0°C. Then the mixture was added slowly to a solution of ammonia (25%, 22 mL) in EtOAc (20 mL) at 0°C (pH ≈ 9). The cooling bath was removed and the mixture was stirred vigorously at rt for 23 h. The mixture was diluted with water and EtOAc and extracted with EtOAc against brine (5x). The combined organic phases were dried over magnesium sulfate and the solvent was removed *in vacuo*. The crude product was purified by preparative HPLC (C18, 5 µm, 50 mL/min, 30 – 60% MeCN (0.1% FA)/$H_2O$ (0.1% FA), $R_t$ = 25.7 min). After lyophilization the pure product **04** was obtained as a colorless solid (264 mg, 44%). $^{1}$H NMR (400 MHz, $CD_3CN$): δ = 7.35 (dd, $J$ = 8.5, 2.3 Hz, 1H),

7.28 (d, *J* = 2.3 Hz, 1H), 6.91 (d, *J* = 8.6 Hz, 1H), 5.55 (s, 2H), 4.86 (s, 2H), 4.19 (s, 4H), 4.13 (q, *J* = 7.1 Hz, 4H), 3.63 (dt, *J* = 9.9, 4.8 Hz, 4H), 3.47 (dt, *J* = 29.0, 4.8 Hz, 4H), 1.21 (t, *J* = 7.1 Hz, 6H). $^{13}C$ NMR (101 MHz, $CD_3CN$): δ 171.7, 166.4, 153.4, 140.0, 136.7, 120.6, 117.5, 113.6, 67.2, 67.1, 66.8, 61.5, 54.2, 45.8, 42.7, 14.5. HRMS ($ESI^+$) m/z calcd. for $C_{20}H_{29}N_3O_9S$ $[M+Na]^+$ 510.1517, found 510.1516.

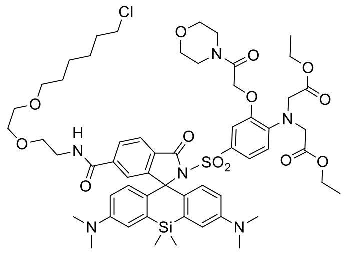

**S05**

SiR-CA (4.30 mg, 6.33 µmol, 1.0 eq.) was dissolved in anhydrous dichloromethane (1.3 mL). Freshly distilled thionyl chloride (6.89 µL, 94.9 µmol, 15 eq.) and anhydrous pyridine (4.30 µL, 53.2 µmol, 8.4 eq.) were added. The reaction mixture was stirred at rt for 30 min before the solvent was evaporated at 50°C until almost anhydrous. In a separate vessel MOBHA-sulfonamide **04** (4.30 mg, 8.82 µmol, 1.4 eq.) was dissolved in anhydrous acetonitrile (2.0 mL), and DIPEA (31.4 µL, 190 µmol, 30 eq.) and DMAP (387 µg, 3.17 µmol, 0.5 eq.) were added. The solution of **04** was heated to 50°C and added to the previously concentrated reaction mixture. After the reaction was stirred at 60°C for 2.5 h the solvent was removed *in vacuo*. The residue was purified by preparative HPLC (8 mL/min, 30 – 80% MeCN/$H_2O$ (0.1% TFA), $R_t$ = 39.3 min) to yield the pure product **S05** as a turquoise solid (1.00 mg, 14%). $^1H$ NMR (400 MHz, DMSO-$d_6$): δ = 9.04 (s, 1H), 8.65 (t, *J* = 5.6 Hz, 1H), 7.89 (s, 2H), 7.06 – 7.01 (m, 2H), 6.91 (d, *J* = 2.9 Hz, 2H), 6.85 (d, *J* = 8.9 Hz, 1H), 6.65 (d, *J* = 2.2 Hz, 1H), 6.50 (dd, *J* = 9.2, 2.8 Hz, 1H), 6.34 (d, *J* = 9.1 Hz, 2H), 4.92 (s, 2H), 4.05 (q, *J* = 7.1 Hz, 4H), 4.00 (s, 4H), 3.61 – 3.55 (m, 4H), 3.57 (t, *J* = 6.6 Hz, 2H), 3.45 – 3.40 (m, 2H), 3.40 – 3.37 (m, 2H), 3.27 (t, *J* = 6.5 Hz, 4H), 3.10 (q, *J* = 7.3 Hz, 2H), 3.09 (q, *J* = 7.3 Hz, 2H), 2.93 (s, 12H), 1.64 (p, *J* = 6.7 Hz, 2H), 1.39 (p, *J* = 7.0 Hz, 2H), 1.34 – 1.27 (m, 2H), 1.26 – 1.21 (m, 2H), 1.15 (t, *J* = 7.1 Hz, 6H), 0.58 (s, 1H), 0.52 (s, 1H). $^{13}C$ NMR (101 MHz, DMSO-$d_6$): δ 170.2, 166.0, 165.2, 164.8, 157.9, 157.6, 155.9, 153.0, 148.2, 140.4, 137.9, 135.5, 134.7, 130.8, 129.9, 129.1, 128.3, 127.0, 122.3, 115.0, 114.3, 78.3, 75.0, 70.1, 69.5, 69.3, 68.6, 66.0, 65.9, 65.7, 60.1, 52.8, 45.7, 45.3, 32.0, 29.0, 26.0, 24.8, 14.1, 8.6, -0.1, -0.7. HRMS ($ESI^+$) m/z calcd. for $C_{57}H_{75}ClN_6O_{13}SSi$ $[M+H]^+$ 1147.4643, found 1147.4645.

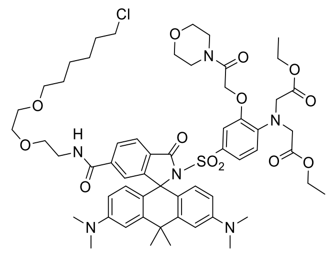

**S06**

CPY-CA (5.00 mg, 7.55 µmol, 1.0 eq.) was dissolved in anhydrous dichloromethane (1.6 mL). Freshly distilled thionyl chloride (8.22 µL, 113 µmol, 15 eq.) and anhydrous pyridine (4.89 µL, 60.4 µmol, 8.0 eq.) were added. The reaction mixture was stirred at rt for 30 min before the solvent was evaporated at 50°C until almost anhydrous. In a separate vessel MOBHA-sulfonamide **04** (5.00 mg, 10.3 µmol, 1.4 eq.) was dissolved in anhydrous acetonitrile (2.0 mL), and DIPEA (31.4 µL, 190 µmol, 30 eq.) and DMAP (387 µg, 3.17 µmol, 0.5 eq.) were added. The solution of **04** was heated to 50°C and added to the previously evaporated reaction mixture. After the reaction was stirred at 60°C for 3.5 h the solvent was removed *in vacuo*. The residue was purified by preparative HPLC (8 mL/min, 30 – 80% MeCN/$H_2O$ (0.1% TFA), $R_t$ = 33.6 min) to yield the pure product **S06** as a blue-turquoise solid (2.30 mg, 27%). $^1$H NMR (400 MHz, DMSO-$d_6$): δ = 9.07 (s, 1H), 8.67 (t, *J* = 5.6 Hz, 1H), 7.89 (s, 2H), 7.14 (dd, *J* = 8.7, 2.2 Hz, 1H), 7.04 – 7.03 (m, 1H), 6.89 (d, *J* = 9.2 Hz, 1H), 6.87 (d, *J* = 2.8 Hz, 1H), 6.64 (d, *J* = 2.2 Hz, 1H), 6.42 (dd, *J* = 8.9, 2.5 Hz, 2H), 6.28 (d, *J* = 8.9 Hz, 2H), 4.94 (s, 2H), 4.03 (q, *J* = 7.1 Hz, 4H), 4.01 (s, 4H), 3.64 – 3.52 (m, 4H), 3.61 – 3.52 (m, 2H), 3.44 – 3.40 (m, 4H), 3.40 – 3.36 (m, 2H), 3.27 (t, *J* = 6.4 Hz, 4H), 3.10 (qd, *J* = 7.3, 4.7 Hz, 4H), 2.95 (s, 12H), 1.85 (d, *J* = 2.2 Hz, 4H), 1.70 – 1.59 (m, 2H), 1.43 – 1.35 (m, 2H), 1.34 – 1.27 (m, 2H), 1.26 – 1.21 (m, 2H), 1.14 (t, *J* = 7.1 Hz, 6H). $^{13}$C NMR (101 MHz, DMSO-$d_6$): δ 170.2, 165.9, 165.2, 164.8, 157.9, 157.6, 155.4, 153.0, 149.6, 145.2, 140.5, 137.9, 130.9, 128.4, 127.8, 127.4, 121.9, 118.0, 116.9, 112.0, 111.7, 109.6, 71.7, 70.1, 69.5, 69.3, 68.6, 66.0, 65.7, 60.1, 52.8, 45.7, 45.3, 44.5, 41.6, 37.6, 35.6, 32.7, 32.0, 29.0, 26.1, 24.9, 21.1, 14.1, 8.6. HRMS (ESI$^+$) m/z calcd. for $C_{58}H_{75}ClN_6O_{13}S$ [M+H]$^+$ 1131.4874, found 1131.4880.

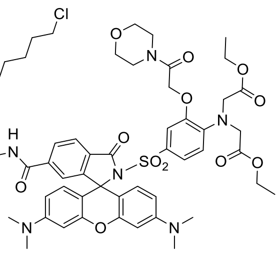

**S07**

TMR-CA (5.40 mg, 8.47 µmol, 1.0 eq.) and MOBHA-sulfonamide **04** (4.96 mg, 10.2 µmol, 1.2 eq.) were dissolved in anhydrous dichloromethane (1.7 mL) in a crimp-top vial. Then DMAP (7.82 mg, 64.0 µmol, 8.0 eq.) and EDC-HCl (12.3 mg, 64.0 µmol, 8.0 eq.) were added. The reaction mixture was then stirred at 65°C for 22 h. After evaporating the solvent *in vacuo,* the residue was purified by preparative HPLC (8 mL/min, 30 – 80% MeCN/$H_2O$ (0.1% TFA), $R_t$ = 34.1 min) to yield the pure product **S07** as a purple solid (3.00 mg, 32%). $^1$H NMR (400 MHz, DMSO-$d_6$): δ = 8.69 (t, *J* = 5.6 Hz, 1H), 8.02 (dd, *J* = 8.1, 1.4 Hz, 1H), 7.90 (d, *J* = 8.0 Hz, 1H), 7.45 – 7.42 (m, 1H), 6.95 (dd, *J* = 8.7, 2.2 Hz, 1H), 6.86 (d, *J* = 8.8 Hz, 1H), 6.79 (d, *J* = 2.2 Hz, 1H), 6.47 (d, *J* = 2.0 Hz, 2H), 6.41 – 6.30 (m, 4H), 4.94 (s, 2H), 4.07 (s, 4H), 4.03 (q, *J* = 7.1 Hz, 4H), 3.63 – 3.52 (m, 8H), 3.58 (t, *J* = 6.6 Hz, 2H), 3.44 – 3.43 (m, 2H), 3.28 (t, *J* = 6.5 Hz, 4H), 3.15 – 3.05 (m, 2H), 2.96 (s, 12H), 1.65 (p, *J* = 6.8 Hz, 2H), 1.39 (p, *J* = 7.0 Hz, 2H), 1.35 – 1.28 (m, 2H), 1.27 – 1.20 (m, 2H), 1.14 (t, *J* = 7.1 Hz, 6H). $^{13}$C NMR (101 MHz, DMSO-$d_6$): δ 170.2, 165.2, 164.8, 164.6, 158.0, 157.7, 153.3, 152.4, 151.2, 140.4, 138.0, 130.8, 129.3, 128.3, 123.7, 122.8, 122.2, 117.3, 112.0, 108.4, 106.2, 98.2, 70.1, 69.5, 69.3, 68.6, 66.0, 65.8, 60.2, 52.9, 45.8, 45.4, 44.5, 41.6, 32.0, 29.0, 26.1, 24.9, 14.1, 8.6. HRMS (ESI$^+$) m/z calcd. for $C_{55}H_{69}ClN_6O_{14}S$ $[M+H]^+$ 1105.4354, found 1105.4352; m/z calcd. for $C_{55}H_{69}ClN_6O_{14}S$ $[M+2H]^{2+}$ 553.2213, found 553.2207.

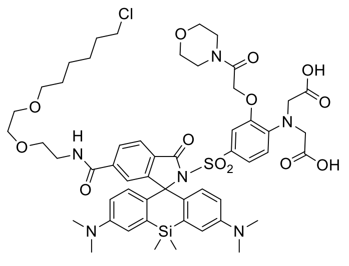

**MaPCa-656$_{low}$**

SiR-CA-MOBHA-COOEt **S05** (2.00 mg, 1.74 µmol, 1.0 eq.) was dissolved in THF (1.0 mL). Then methanol (1.0 mL) and potassium hydroxide solution (0.1M, 1.3 mL) were added. The reaction mixture was stirred at rt for 6 h. The reaction was neutralized by addition of hydrochloric acid (0.1M, 0.2 mL) and acetic acid (≥ 99%, 5.0 µL) before the solvent was evaporated *in vacuo*. The residue was purified by preparative HPLC (8 mL/min, 30 – 70% MeCN/$H_2O$ (0.1% TFA), $R_t$ = 30.6 min) to yield the pure product **MaPCa-656$_{low}$** as a light green solid (1.10 mg, 58%). $^1$H NMR (400 MHz, DMSO-d$_6$): δ = 12.42 (bs, 2H), 9.04 (s, 1H), 8.65 (t, *J* = 5.6 Hz, 1H), 7.94 – 7.84 (m, 2H), 7.04 (s, 1H), 6.96 (dd, *J* = 8.7, 2.2 Hz, 1H), 6.91 (d, *J* = 2.9 Hz, 1H), 6.82 (d, *J* = 8.9 Hz, 1H), 6.75 (d, *J* = 2.2 Hz, 1H), 6.50 (dd, *J* = 9.1, 2.8 Hz, 2H), 6.34 (d, *J* = 9.0 Hz, 2H), 4.92 (s, 2H), 3.93 (s, 4H), 3.57 (t, *J* = 6.6 Hz, 2H), 3.41 – 3.37 (m, 2H), 3.27 (t, *J* = 6.5 Hz, 4H), 3.10 (qd, *J* = 7.3, 4.7 Hz, 4H), 2.92 (s, 12H), 1.65 (p, *J* = 6.8 Hz, 2H), 1.39 (p, *J* = 6.8 Hz, 2H), 1.34 – 1.27 (m, 2H), 1.27 – 1.20 (m, 2H), 0.58 (s, 1H), 0.54 (s, 1H). $^{13}$C NMR (101 MHz, DMSO-d$_6$): δ 171.7, 166.1, 165.3, 164.8, 162.5, 158.1, 157.7, 155.9, 153.0, 148.1, 147.0, 140.4, 139.6, 130.8, 129.2, 128.4, 127.0, 126.7, 121.5, 115.1, 114.3, 113.0, 75.1, 70.1, 69.5, 69.3, 68.6, 65.9, 65.7, 52.5, 45.7, 45.3, 32.0, 29.0, 26.1, 24.9, 8.6, -0.1, -0.7. HRMS (ESI$^+$) m/z calcd. for $C_{53}H_{67}ClN_6O_{13}SSi$ $[M+2H]^{2+}$ 546.2045, found 546.2040.

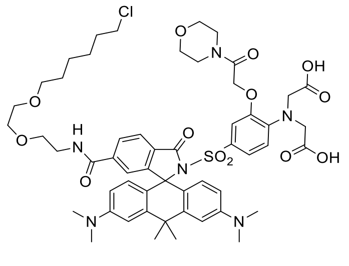

**MaPCa-619$_{low}$**

CPY-CA-MOBHA-COOEt **S06** (5.40 mg, 4.77 µmol, 1.0 eq.) was dissolved in THF (1.0 mL). Then methanol (1.0 mL) and potassium hydroxide solution (0.1M, 1.3 mL) were added. The reaction mixture was stirred at rt for 4 h. The reaction was neutralized by addition of hydrochloric acid (0.1M, 0.2 mL) and acetic acid (≥ 99%, 6.0 µL) before the solvent was evaporated *in vacuo*. The residue was purified by preparative HPLC (8 mL/min, 30 – 70% MeCN/$H_2O$ (0.1% TFA),

$R_t$ = 33.0 min) to yield the pure product **MaPCa-619$_{low}$** as a light blue solid (1.20 mg, 23%). $^{1}$H NMR (400 MHz, DMSO-$d_6$): δ = 12.41 (s, 2H), 8.67 (t, *J* = 5.7 Hz, 1H), 7.90 (s, 2H), 7.07 (dd, *J* = 8.7, 2.2 Hz, 1H), 7.03 (s, 1H), 6.89 (d, *J* = 2.6 Hz, 1H), 6.86 (d, *J* = 8.9 Hz, 1H), 6.75 (d, *J* = 2.3 Hz, 1H), 6.45 (dd, *J* = 9.0, 2.6 Hz, 2H), 6.30 (d, *J* = 8.8 Hz, 2H), 4.94 (s, 2H), 3.95 (s, 4H), 3.57 (t, *J* = 6.6 Hz, 2H), 3.27 (t, *J* = 6.5 Hz, 2H), 3.10 (qd, *J* = 7.3, 4.8 Hz, 2H), 2.95 (s, 12H), 1.85 (d, *J* = 4.4 Hz, 6H), 1.65 (p, *J* = 6.7 Hz, 2H), 1.39 (p, *J* = 7.0 Hz, 2H), 1.34 – 1.27 (m, 2H), 1.26 – 1.20 (m, 2H). $^{13}$C NMR (101 MHz, DMSO-$d_6$): δ 171.8, 165.9, 165.3, 164.9, 158.6, 158.1, 153.0, 149.5, 145.2, 140.4, 138.4, 130.9, 128.4, 127.8, 127.3, 123.8, 117.2, 112.0, 111.9, 71.7, 70.1, 69.5, 69.4, 68.6, 66.0, 52.4, 45.8, 45.4, 44.6, 41.6, 37.6, 35.7, 32.8, 32.0, 29.0, 26.1, 24.9, 8.6. HRMS (ESI$^+$) m/z calcd. for $C_{54}H_{67}ClN_6O_{13}S$ [M+2H]$^{2+}$ 538.2160, found 538.2157.

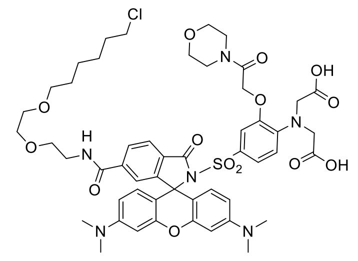

**MaPCa-558$_{low}$**

TMR-CA-MOBHA-COOEt **S07** (4.40 mg, 3.98 µmol, 1.0 eq.) was dissolved in THF (1.0 mL). Then methanol (1.0 mL) and potassium hydroxide solution (0.1M, 1.3 mL) were added. The reaction mixture was stirred at rt for 5 h. The reaction was neutralized by addition of hydrochloric acid (0.1M, 0.2 mL) and acetic acid (≥ 99%, 5.0 µL) before the solvent was evaporated *in vacuo*. The residue was purified by preparative HPLC (8 mL/min, 30 – 70% MeCN/$H_2O$ (0.1% TFA), $R_t$ = 30.3 min) to yield the pure product **MaPCa-558$_{low}$** as a dark purple solid (0.90 mg, 22%). $^{1}$H NMR (400 MHz, DMSO-$d_6$): δ = 12.44 (bs, 2H), 8.69 (t, *J* = 5.6 Hz, 1H), 8.02 (d, *J* = 8.1 Hz, 1H), 7.91 (d, *J*= 8.0 Hz, 1H), 7.42 (s, 1H), 7.02 (dd, *J* = 8.7, 2.2 Hz, 1H), 6.86 (d, *J* = 8.9 Hz, 1H), 6.74 (d, *J* = 2.3 Hz, 1H), 6.46 (d, *J* = 2.2 Hz, 2H), 6.41 – 6.28 (m, 4H), 4.94 (s, 2H), 3.97 (s, 4H), 3.58 (t, *J* = 6.6 Hz, 2H), 3.31 – 3.25 (m, 2H), 3.14 – 3.06 (m, 2H), 2.95 (s, 12H), 1.65 (p, *J* = 6.7 Hz, 2H), 1.40 (p, *J* = 7.0 Hz, 2H), 1.35 – 1.27 (m, 2H), 1.27 – 1.19 (m, 2H). $^{13}$C NMR (101 MHz, DMSO-$d_6$): δ 171.7, 165.3, 164.8, 164.6, 157.8, 153.3, 153.2, 152.3, 151.2, 140.3, 138.4, 130.7, 129.3, 128.4, 117.1, 114.5, 110.9, 108.4, 106.2, 98.2, 70.1, 69.5, 69.3, 68.6, 66.0, 65.7, 52.5, 45.4, 32.0, 29.0, 26.1, 24.9, 5.4. HRMS (ESI$^+$) m/z calcd. for $C_{51}H_{61}ClN_6O_{14}S$ [M+H]$^+$ 1049.3728, found 1049.3729; m/z calcd. for $C_{51}H_{61}ClN_6O_{14}S$ [M+2H]$^{2+}$ 525.1900, found 525.1896.

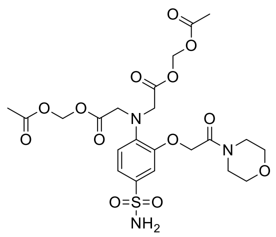

**06**

MOBHA-sulfonamide **05** (50.0 mg, 103 µmol, 1.0 eq.) was dissolved in acetonitrile (9.0 mL). Then DMAP (6.26 mg, 51.3 µmol, 0.5 eq.) and di-tert-butyl-carbonate (110 µL, 513 µmol, 5.0 eq.) in acetonitrile (1.0 mL) was added. The reaction was stirred at rt for 19 h followed by evaporation of the solvent *in vacuo.* di-tert-butyl-carbonate (110 µL, 513 µmol, 5.0 eq.) and DMAP (6.26 mg, 51.3 µmol, 0.5 eq.) were dissolved in DCM (8.0 mL) and added to the residue. The reaction mixture was stirred at 35°C for 24 h. Afterwards the reaction was quenched by addition of water and extracted with DCM (3x). The combined organic phases were dried over magnesium sulfate and the solvent was removed *in vacuo*. The crude product was purified by flash column chromatography (12 g $SiO_2$, 0 – 5% MeOH/DCM) to yield the intermediate product as a beige-yellow solid (52.4 mg, 79%). It was directly re-dissolved in THF (2.0 mL). Then methanol (2.0 mL) and potassium hydroxide solution (0.1M, 2.8 mL) were added. The reaction mixture was stirred at rt for 5.5 h followed by neutralizing with hydrochloric acid (0.1M, 0.4 mL) and acetic acid (≥ 99%, 2.8 µL) and evaporation of the solvents *in vacuo*. The residue was suspended in acetonitrile (2.0 mL). Bromomethyl acetate (73.7 µL, 752 µmol, 8.8 eq.) and DIPEA (355 µL, 2.15 mmol, 25 eq.) were added. After stirring the reaction at rt for 21 h the solvent was evaporated *in vacuo* again. The residue was suspended in DCM (0.9 mL) and triisopropyl silane (135 µL, 7.5% v/v) and TFA (0.9 mL) were added. The reaction was stirred at rt for 30 min and then neutralized by addition of a saturated $NaHCO_3$-solution (1.2 mL). Extraction with EtOAc (5x) against brine, drying of the combined organic phases over magnesium sulfate and evaporation of the solvent *in vacuo* led to the crude product. After purification *via* preparative HPLC (C18, 5 µm, 50 mL/min, 30 – 50% MeCN (0.1% FA)/$H_2O$ (0.1% FA), $R_t$ = 22.9 min) the pure product **06** was obtained as a colorless solid (22.4 mg, 36% over 5 steps). $^1H$ NMR (400 MHz, $CD_3CN$): δ = 7.38 (dd, *J* = 8.6, 2.2 Hz, 1H), 7.23 (d, *J* = 2.2 Hz, 1H), 6.92 (d, *J* = 8.6 Hz, 1H), 5.71 (s, 4H), 5.58 (s, 2H), 4.88 (s, 2H), 4.26 (s, 4H), 3.63 (dt, *J* = 12.7, 4.8 Hz, 4H), 3.47 (dt, *J* = 29.6, 4.8 Hz, 4H), 2.06 (s, 6H). $^{13}C$ NMR (101 MHz, $CD_3CN$) δ 170.7, 170.7, 166.4, 153.4, 139.4, 136.8, 121.0, 117.4, 113.6, 80.2, 66.7, 53.9, 45.7, 42.7, 20.9. HRMS ($ESI^+$) m/z calcd. for $C_{22}H_{29}N_3O_{13}S$ $[M+H]^+$ 576.1494, found 576.1495.

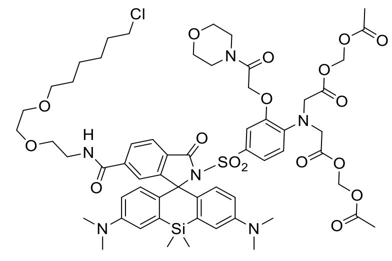

**MaPCa-656$_{low}$ AM**

SiR-CA (5.00 mg, 7.36 µmol, 1.0 eq.) was dissolved in DCM (1.6 mL). Freshly distilled thionyl chloride (8.01 µL, 110 µmol, 15 eq.) and anhydrous pyridine (4.76 µL, 58.9 µmol, 8.0 eq.) were added. The reaction mixture was stirred at rt for 30 min before the solvent was evaporated at 50°C until almost dry. In a separate vessel MOBHA-AM-sulfonamide **06** (5.51 mg, 9.57 µmol, 1.3 eq.) was dissolved in anhydrous acetonitrile (2.3 mL), and DIPEA (36.5 µL, 221 µmol, 30 eq.) and DMAP (450 µg, 3.68 µmol, 0.5 eq.) were added. The solution of **06** was heated to 50°C and added to the previously evaporated reaction mixture. After the reaction was stirred at 60°C for 3 h the solvent was removed *in vacuo*. The residue was purified by preparative HPLC (8 mL/min, 30 – 70% MeCN/$H_2O$ (0.1% TFA), $R_t$ = 40.2 min) to yield the pure product **MaPCa-656$_{low}$ AM** as a turquoise solid (2.20 mg, 24%). $^1$H NMR (400 MHz, DMSO-$d_6$): δ = 8.66 (t, *J* = 5.6 Hz, 1H), 7.92 (d, *J* = 8.1 Hz, 1H), 7.88 (dd, *J* = 8.1, 1.4 Hz, 1H), 7.05 – 7.04 (m, 1H), 6.97 (dd, *J* = 8.8, 2.2 Hz, 1H), 6.93 (d, *J* = 2.9 Hz, 2H), 6.85 (d, *J* = 8.9 Hz, 1H), 6.63 (d, *J* = 2.2 Hz, 1H), 6.50 (dd, *J* = 9.1, 2.9 Hz, 2H), 6.32 (d, *J* = 9.0 Hz, 2H), 5.69 (s, 4H), 4.92 (s, 2H), 4.09 (s, 4H), 3.57 (t, *J* = 6.6 Hz, 2H), 3.47 – 3.34 (m, 8H), 3.47 – 3.35 (m, 4H), 3.27 (t, *J* = 6.4 Hz, 4H), 2.93 (s, 12H), 2.04 (s, 6H), 1.64 (p, *J* = 6.8 Hz, 2H), 1.39 (p, *J* = 7.0 Hz, 2H), 1.35 – 1.25 (m, 2H), 1.28 – 1.17 (m, 2H), 0.59 (s, 3H), 0.53 (s, 3H). $^{13}$C NMR (101 MHz, DMSO-$d_6$): δ 169.3, 169.3, 166.1, 165.2, 164.8, 158.3, 158.0, 155.9, 152.9, 148.1, 140.5, 137.4, 134.9, 130.8, 129.2, 128.4, 127.1, 124.0, 122.3, 121.8, 116.5, 115.2, 114.4, 112.1, 79.2, 75.0, 70.1, 69.5, 69.4, 68.6, 66.0, 65.9, 65.6, 52.5, 45.4, 44.5, 41.6, 32.0, 29.0, 26.1, 24.9, 20.5, -0.2, -0.8. HRMS (ESI$^+$) m/z calcd. for $C_{59}H_{75}ClN_6O_{17}SSi$ $[M+H]^+$ 1235.4440, found 1235.4432; m/z calcd. for $C_{59}H_{75}ClN_6O_{17}SSi$ $[M+2H]^{2+}$ 618.2257, found 618.2263.

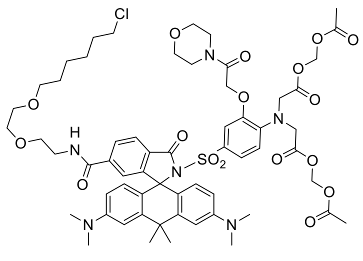

**MaPCa-619$_{low}$ AM**

CPY-CA (5.00 mg, 7.55 µmol, 1.0 eq.) was dissolved in DCM (1.6 mL). Freshly distilled thionyl chloride (8.22 µL, 112 µmol, 15 eq.) and anhydrous pyridine (4.89 µL, 60.4 µmol, 8.0 eq.) were added. The reaction mixture was stirred at rt for 30 min before the solvent was evaporated at 50°C until almost dry. In a separate vessel MOBHA-AM-sulfonamide **06** (5.43 mg, 9.44 µmol, 1.3 eq.) was dissolved in anhydrous acetonitrile (2.4 mL), and DIPEA (37.4 µL, 227 µmol, 30 eq.) and DMAP (461 µg, 3.78 µmol, 0.5 eq.) were added. The solution of **06** was heated to 50°C and added to the previously evaporated reaction mixture. After the reaction was stirred at 60°C for 2.5 h the solvent was removed *in vacuo*. The residue was purified by preparative HPLC (8 mL/min, 30 – 70% MeCN/$H_2O$ (0.1% TFA), $R_t$ = 35.1 min) to yield the pure product **MaPCa-619$_{low}$ AM** as a dark blue solid (1.40 mg, 15%). $^1$H NMR (400 MHz, DMSO-d$_6$): δ = 8.68 (t, *J* = 5.7 Hz, 1H), 7.94 – 7.87 (m, 2H), 7.08 (dd, *J* = 8.6, 2.2 Hz, 1H), 7.04 (s, 1H), 6.91 (s, 2H), 6.88 (s, 1H), 6.60 (d, *J* = 2.2 Hz, 1H), 6.42 (d, *J* = 8.9 Hz, 2H), 6.26 (d, *J* = 8.8 Hz, 2H), 5.68 (s, 4H), 4.94 (s, 2H), 4.10 (s, 4H), 3.57 (t, *J* = 6.6 Hz, 2H), 3.48 – 3.36 (m, 4H), 3.47 – 3.34 (m, 8H), 3.27 (t, *J* = 6.2 Hz, 4H), 2.95 (s, 12H), 2.04 (s, 6H), 1.85 (s, 6H), 1.65 (p, *J* = 6.8 Hz, 2H), 1.39 (p, *J* = 7.1 Hz, 2H), 1.35 – 1.27 (m, 2H), 1.27 – 1.19 (m, 2H). $^{13}$C NMR (101 MHz, DMSO-d$_6$): δ 169.3, 169.2, 166.0, 165.2, 164.8, 158.2, 157.9, 155.3, 152.9, 149.4, 145.2, 140.5, 137.4, 130.9, 128.4, 127.8, 127.4, 123.8, 122.4, 121.7, 116.5, 112.2, 111.9, 109.8, 79.1, 71.6, 70.1, 69.5, 69.3, 68.6, 66.0, 65.9, 65.6, 52.5, 45.3, 44.4, 41.6, 37.6, 35.6, 32.7, 32.0, 29.0, 26.1, 24.9, 20.5, 8.6. HRMS (ESI$^+$) m/z calcd. for $C_{60}H_{75}ClN_6O_{17}S$ [M+H]$^+$ 1219.4671, found 1219.4670; m/z calcd. for $C_{60}H_{75}ClN_6O_{17}S$ [M+2H]$^{2+}$ 610.2372, found 610.2383.

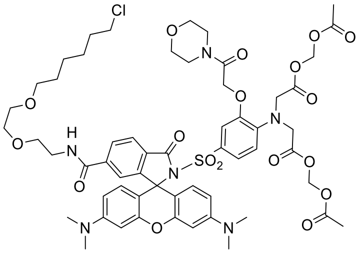

**MaPCa-558$_{low}$ AM**

TMR-CA (5.00 mg, 7.86 µmol, 1.0 eq.) was dissolved in DCM (1.6 mL). Freshly distilled thionyl chloride (8.55 µL, 118 µmol, 15 eq.) and anhydrous pyridine (5.09 µL, 62.9 µmol, 8.0 eq.) were added. The reaction mixture was stirred at rt for 30 min before the solvent was evaporated at 50°C until almost dry. In a separate vessel MOBHA-AM-sulfonamide **06** (5.65 mg, 9.82 µmol, 1.3 eq.) was dissolved in anhydrous acetonitrile (2.5 mL), and DIPEA (39.0 µL, 236 µmol, 30 eq.) and DMAP (480 µg, 3.93 µmol, 0.5 eq.) were added. The solution of **06** was heated to 50°C and added to the previously evaporated reaction mixture. After the reaction was stirred at 60°C for 2.5 h the solvent was removed *in vacuo*. The residue was purified by preparative HPLC (8 mL/min, 30 – 80% MeCN/$H_2O$ (0.1% TFA), $R_t$ = 33.3 min) to yield the pure product **MaPCa-558$_{low}$ AM** as a purple solid (1.20 mg, 13%). $^1$H NMR (400 MHz, DMSO-$d_6$): δ = 8.69 (t, *J* = 5.7 Hz, 1H), 8.02 (dd, *J* = 8.0, 1.4 Hz, 1H), 7.92 (d, *J* = 8.0 Hz, 1H), 7.43 (d, *J* = 1.4 Hz, 1H), 7.01 (dd, *J* = 8.7, 2.2 Hz, 1H), 6.89 (d, *J* = 8.9 Hz, 1H), 6.63 (d, *J* = 2.3 Hz, 1H), 6.46 (d, *J* = 2.4 Hz, 2H), 6.35 (dd, *J* = 9.0, 2.4 Hz, 2H), 6.31 (d, *J* = 8.8 Hz, 2H), 5.68 (s, 4H), 4.94 (s, 2H), 4.14 (s, 4H), 3.58 (t, *J* = 6.6 Hz, 2H), 3.47 – 3.37 (m, 4H), 3.32 – 3.25 (m, 4H), 2.96 (s, 12H), 2.05 (s, 6H), 1.65 (p, *J* = 6.7 Hz, 2H), 1.40 (p, *J* = 7.1 Hz, 2H), 1.35 – 1.28 (m, 2H), 1.27 – 1.19 (m, 2H). $^{13}$C NMR (101 MHz, DMSO-$d_6$): δ 169.3, 169.2, 165.2, 164.8, 164.5, 158.0, 157.7, 153.2, 153.1, 152.4, 151.2, 140.4, 137.4, 130.7, 129.3, 128.3, 123.7, 122.8, 122.2, 116.5, 112.1, 108.3, 106.1, 98.1, 79.0, 70.1, 69.5, 69.3, 68.6, 66.0, 65.9, 65.6, 52.5, 45.3, 44.5, 41.6, 32.0, 29.0, 26.1, 24.9, 20.5. HRMS (ESI$^+$) m/z calcd. for $C_{57}H_{69}ClN_6O_{18}S$ $[M+H]^+$ 1193.4150, found 1193.4145; m/z calcd. for $C_{57}H_{69}ClN_6O_{18}S$ $[M+2H]^{2+}$ 597.2112, found 597.2104.
